# Multi-timescale neural adaptation underlying long-term musculoskeletal reorganization

**DOI:** 10.1101/2024.10.18.618983

**Authors:** Roland Philipp, Yuki Hara, Naohito Ohta, Naoki Uchida, Tomomichi Oya, Tetsuro Funato, Kazuhiko Seki

## Abstract

The central nervous system (CNS) can effectively control body movements despite environmental changes. While much is known about adaptation to external environmental changes, less is known about responses to internal bodily changes. This study investigates how the CNS adapts to long-term alterations in the musculoskeletal system using a tendon transfer model in non-human primates. We surgically relocated finger flexor and extensor muscles to examine how the CNS adapts its strategy for finger movement control by measuring muscle activities during grasping tasks. Two months post-surgery, the monkeys demonstrated significant recovery of grasping function despite the initial disruption. Our findings suggest a two-phase CNS adaptation process: an initial phase enabling function with the transferred muscles, followed by a later phase abandoning this enabled function and restoring a control strategy that, while potentially less conflicted than the maladaptive state, resembled the original pattern, possibly representing a ‘good enough’ solution. These results highlight a multi-phase CNS adaptation process with distinct time constants in response to sudden bodily changes, offering potential insights into understanding and treating movement disorders.

**SIGNIFICANCE STATEMENT:** After major changes to the body’s mechanics, the nervous system adapts using strategies on multiple timescales. Our primate tendon transfer study shows that core muscle synergy groupings remain stable, reflecting a default to modular control. However, the activation of these synergies changes dramatically; an initial, rapid ‘swap’ of their timing proves to be **maladaptive**, impairing motor function. This conflict is only resolved through the gradual development of **slower, compensatory strategies** over several weeks. This process highlights the fundamental tension the CNS faces when its reliance on stable motor modules conflicts with the need for flexible control, offering insights into neural plasticity and staged rehabilitation.

## Introduction

The central nervous system (CNS) continuously adapts bodily functions in response to both external and internal challenges. Experimental models based on external perturbations, such as altered gravitational fields or distorted sensory feedback, have illuminated mechanisms of sensorimotor adaptation (Sugita, 1996; Davidson and Wolpert, 2003; Luauté et al., 2009; Fleury et al., 2019). Because the changes in the external environment can be controlled by the experimental design, these models provide an opportunity to assess how the CNS adapts to them. However, the transient and predictable nature of these changes may not fully capture the demands posed by internal, long-lasting, and unpredictable alterations to the body’s internal environment

In contrast, the internal changes such as developmental growth (Power and Schlaggar, 2017), fatigue (Green, 1997), postural sway (Zemková, 2022), and sensory disorientation (Schärli et al., 2024) impose distinct challenges. Because identifying the source, extent, and time-constant of changes in the internal environment is usually difficult, assessing the corresponding CNS adaptation is also challenging. Particularly, structural alterations to the musculoskeletal system, whether due to injury, disease, or surgery, fundamentally change the body’s biomechanics and sensorimotor associations, but the quantification of these changes is usually difficult. Accordingly, the way CNS remaps its motor control strategies corresponding to the changes is not yet well understood (Walker et al., 2004), although understanding such adaptations is crucial for elucidating the pathophysiology of motor impairments observed in chronic musculoskeletal conditions such as osteoarthritis, a degenerative joint disease, (Hunter and Eckstein, 2009) and muscular dystrophy, which involves progressive muscle weakness, (Mercuri and Muntoni, 2013).

To address this question, in this study, we employed a tendon transfer (TT) surgery model (Sperry, 1940), which introduces a controlled, sustained change to the musculoskeletal structure. TT surgery is a clinically well-established procedure (Gardenier et al., 2020) that surgically re-attaches the tendon of a specific muscle to that of a surrounding one. This procedure relocates a specific muscle, so that its contraction generates a new mechanical action and, consequently, novel somatosensory feedback. Because the internal change is controlled by the experimenter, the TT model provides a powerful platform to investigate how the CNS adapts to a new internal state. Another unique feature of TT model is that it places permanent changes of the internal environment while leaving the CNS anatomically intact. Unlike CNS lesion models (Hoffmann et al., 2009), where the injury itself disrupts neural circuits and thereby complicates the assessment of adaptive capacity of the CNS, TT offers a distinct advantage: it allows investigation of CNS-driven adaptation without the confounding effects of direct neural damage. Tendon transfer has been used in various species, with studies suggesting different adaptive capacities, ranging from limited adaptation in adult rodents (Sperry, 1940; Sperry, 1942; Slawinska and Kasicki, 2002; Bowlus et al., 2003) to more substantial adjustments in cats (Yumiya et al., 1979; Loeb, 1999) and primates, including humans (Lee and Seung, 1999; Wester et al., 2018; Gaetz et al., 2023). Therefore, adaptability to TT appears enhanced in primates and humans. Altogether, this approach provides a controlled platform to examine how the CNS adapts to musculoskeletal changes in primates.

According to earlier reports, there are two motor control strategies that could be involved when the CNS needs to face the sensorimotor remapping posed by structural alterations to the musculoskeletal system. First, the CNS may employ modular building blocks such as muscle synergies, coordinated activations of muscle groups that reduce the dimensionality of motor control (d’Avella et al., 2003; Bizzi and Cheung, 2013). Second, skilled behaviors like fine finger movements often require fractionation, the capacity to activate muscles independently. When confronted with structural changes to the musculoskeletal system, does the CNS adapt by modulating existing synergies, or by shifting toward more fractionated control strategies? Addressing this question requires examining the evolution of neural control strategies over extended periods of recovery to capture both immediate adjustments and long-term learning.

This study aimed to identify long-term adaptive mechanisms of the primate CNS following structural changes to the musculoskeletal system. We investigated whether the CNS adapts by modulating existing muscle synergies or by altering the fractionated control of muscles specifically affected by the surgical alteration. We surgically altered the limb structure by performing crossed tendon transfers of finger extensor and flexor muscle tendons. Using a trained finger-grasping task as our behavioral readout, we examined how the CNS recalibrates muscle activity to regain skilled motor function. Our findings provide new insights into the dynamic reorganization of motor control following structural changes of the body.

## RESULTS

In this study, we developed a novel crossed tendon transfer animal model. This involved surgically swapping the tendons of two antagonistic finger muscles in a macaque monkeys’ forearm (Fig. 1; A complete list of all muscle abbreviations, their full names, and their assigned synergies for each monkey is provided in Supplementary Table 1). The effectiveness and consistency of the surgery was confirmed by measuring: (i) the distance traveled by muscle fibers and their intramuscular tendons in the transferred muscle (Fig. 2B, C); and (ii) the amount of fingertip displacement (Fig. 2D, E). Both measurements were induced by percutaneous electrical stimulation over the transferred muscles (Fig. 2A). For a detailed description of the results of these measurements see the Methods section. The two monkeys were assigned slightly different tasks, allowing us to examine controlled grasping in Monkey A and a more natural grasp in Monkey B (refer to the Methods section, and Figs. 3 and Fig. 4 for further details). The two monkeys ultimately performed related but distinct grasping tasks, a methodological divergence that provided a valuable opportunity to test the generality of the core adaptive mechanisms. Monkey A performed a controlled grasping task requiring a fine precision grip, designed to study adaptation of fine motor control (Fig. 3). While the same task was initially planned for both animals, Monkey B performed this task inconsistently, frequently varying its grip strategy, and we could not reinforce this monkey to perform in single strategy consistently. Therefore, to ensure reliable task engagement and in accordance with the ethical principle of Reduction and Refinement (the ‘3Rs’; (Tannenbaum and Bennett, 2015)), the task for Monkey B was modified to a more naturalistic food retrieval grasp that the animal performed consistently (Fig. 4).

**Figure 1:**
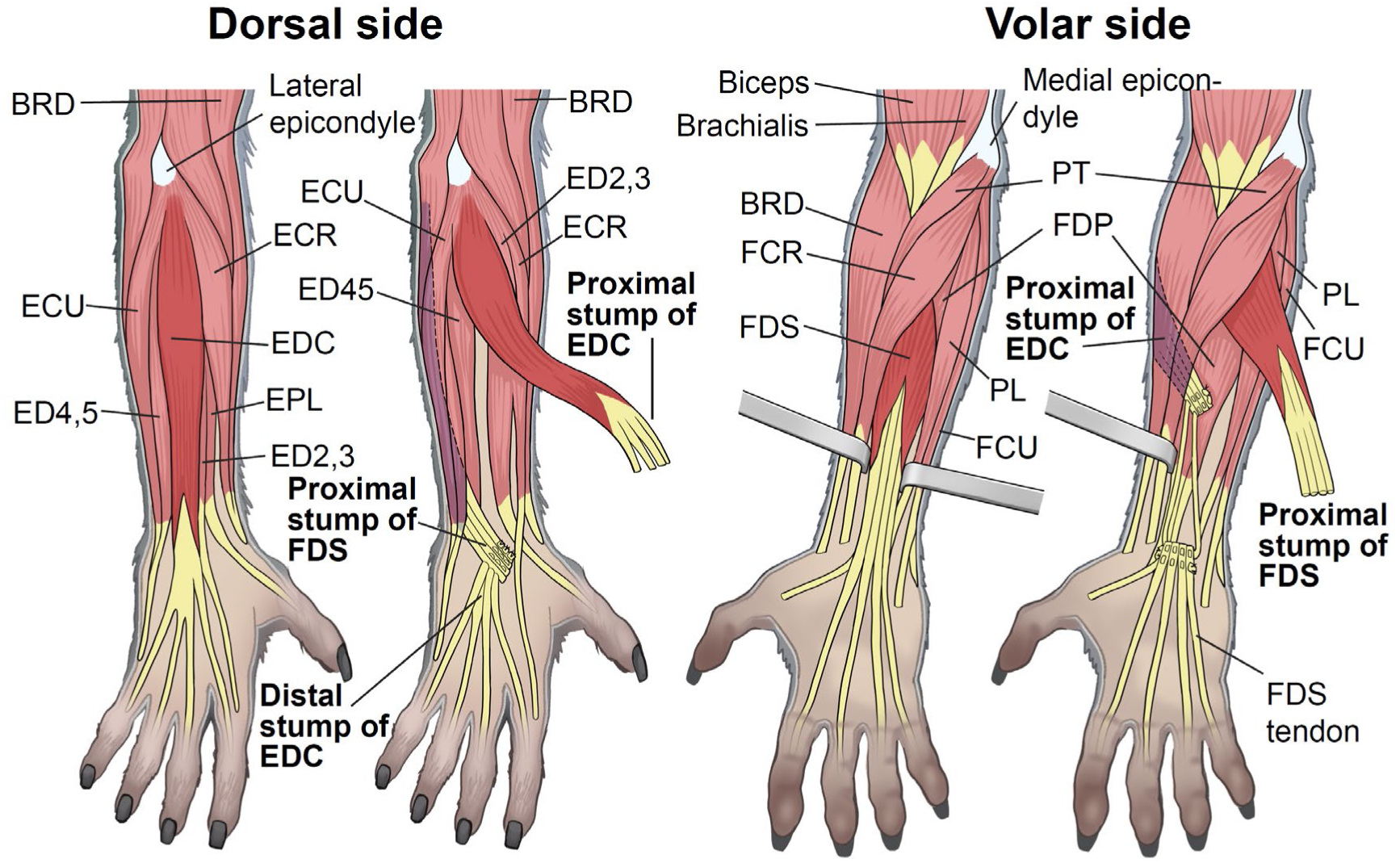
Muscle anatomy of the macaque forearm and the tendon transfer procedure. A schematic of the primary forearm muscles involved in the study, showing both the dorsal and volar views. The diagram illustrates the surgical crossed tendon transfer of the extensor digitorum communis (EDC) and flexor digitorum superficialis (FDS) tendons. All labeled muscles were implanted with EMG electrodes. Muscle Abbreviations: BRD: brachioradialis, ECR: extensor carpi radialis, ECU: extensor carpi ulnaris, ED2,3: extensor digitorum-2,3, ED4,5: extensor digitorum-4,5, EPL: extensor pollicis longus (not implanted), FCR: flexor carpi radialis, FCU: flexor carpi ulnaris, FDP: flexor digitorum profundus, PL: palmaris longus, PT: pronator teres (not implanted), *(The deltoid (DEL) muscle was also implanted in Monkey B but is not shown as it is a shoulder muscle). See also Supplementary Table 1 for a complete list of all muscle abbreviations, their full names, and their assigned synergies*.

**Figure 2:**
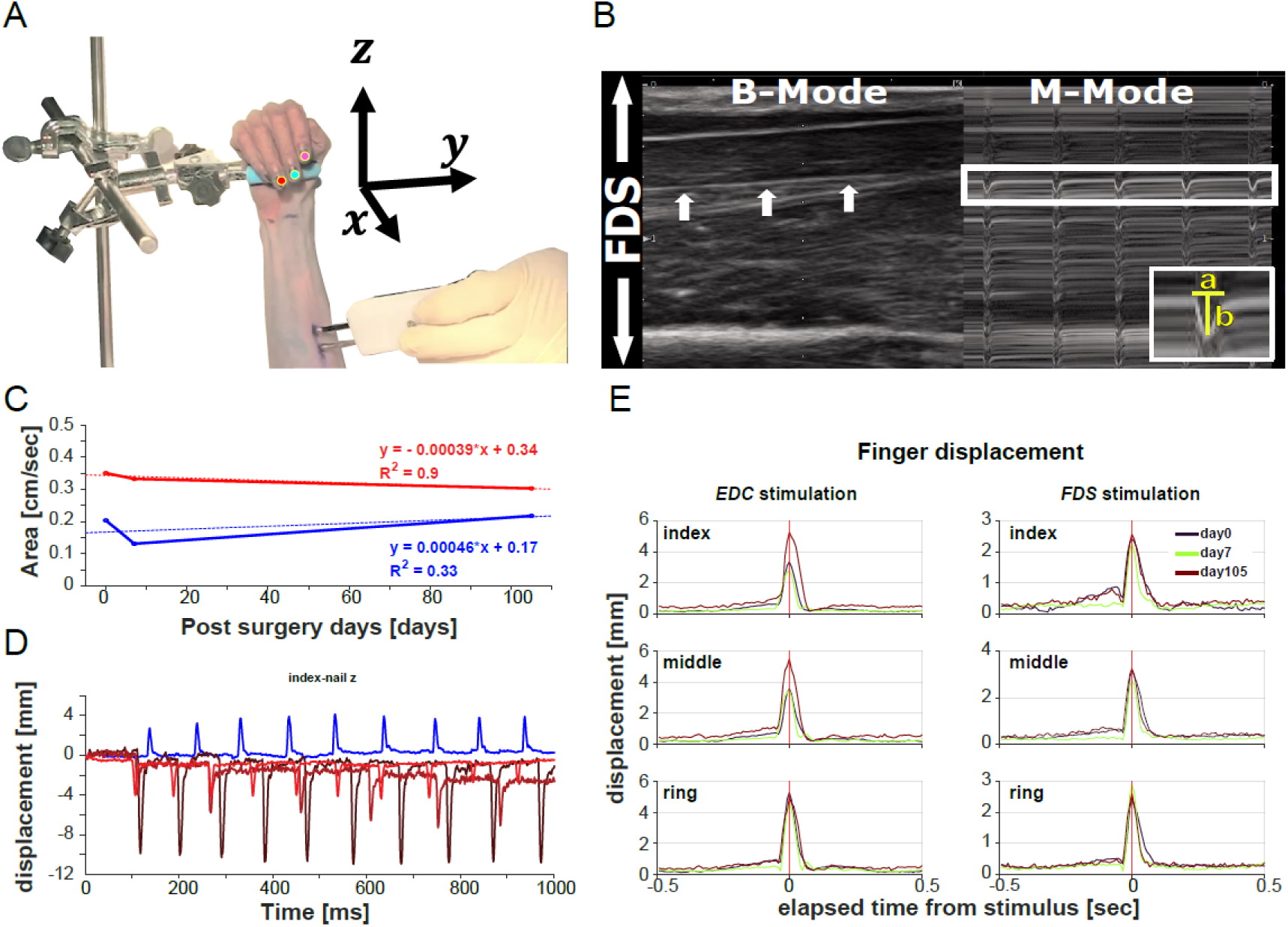
Long term confirmation of tendon surgery effectiveness to alter mechanical properties. (A) Set-up for the ultrasound measurement and video recordings of the stimulation induced movements of the EDC and FDS tendons. (B) Sonogram of the FDS muscle and its intramuscular tendons. Left side (B-mode, i.e., brightness mode) shows the still image of the monkeys forearm at a given point in time. Right side (M-mode, i.e., motion mode) shows the stagged images of the FDS tendon displacement induced by muscle stimulation (50mA). White arrows demarcate the FDS tendon which was used for the measurement. Grayscale gradations correspond to tissue densities: hyperechoic regions (white) denote denser structures like surface of bones and tendons, while hypoechoic areas (black) signify less dense tissues such as adipose tissue and musculature. Inset demonstrates the area measurement. The area of the displacement waves was measured in the M-mode, representing the strength of muscle contraction. We measured the duration (a, sec) and amplitude (b, cm) of three waves and calculated the average. Area = a*b/2(cm/sec) for days 0, 7 and 105 after tendon transfer. (C) Areas under the wave measured in the M-mode for 3 experimental days (0, 7 and 105 days post-TT) and regression lines in red and blue for FDS and EDC, respectively. R^2^>0.5 for FDS. The data suggested that muscle contractions induced by direct electrical stimulation were nearly constant. (D) Markers placed on the index, middle and ring finger nails (A) were used to measure finger displacement in xyz-dimensions. We calculated the sum of the Euclidean distances of each marker from the origin of the 3D coordinate system as a scalar quantity. Observing the movement along the Z-axis, it became reversed post-surgery indicating a reversal from finger flexion to extension due to tendon transfer (D, blue = pre-TT at surgery day; dark brown = post-TT at surgery day; light brown = 1wk post-TT; red = 3wks post-TT). The scalar quantity of the fingers during muscle stimulation did not change much at day 0, 7 and 105 days (E), suggesting that there was no tendon rupture or slackening of the tendons postoperatively (EDC stimulation, left; FDS stimulation, right). Data were collected in monkey A.

**Figure 3:**
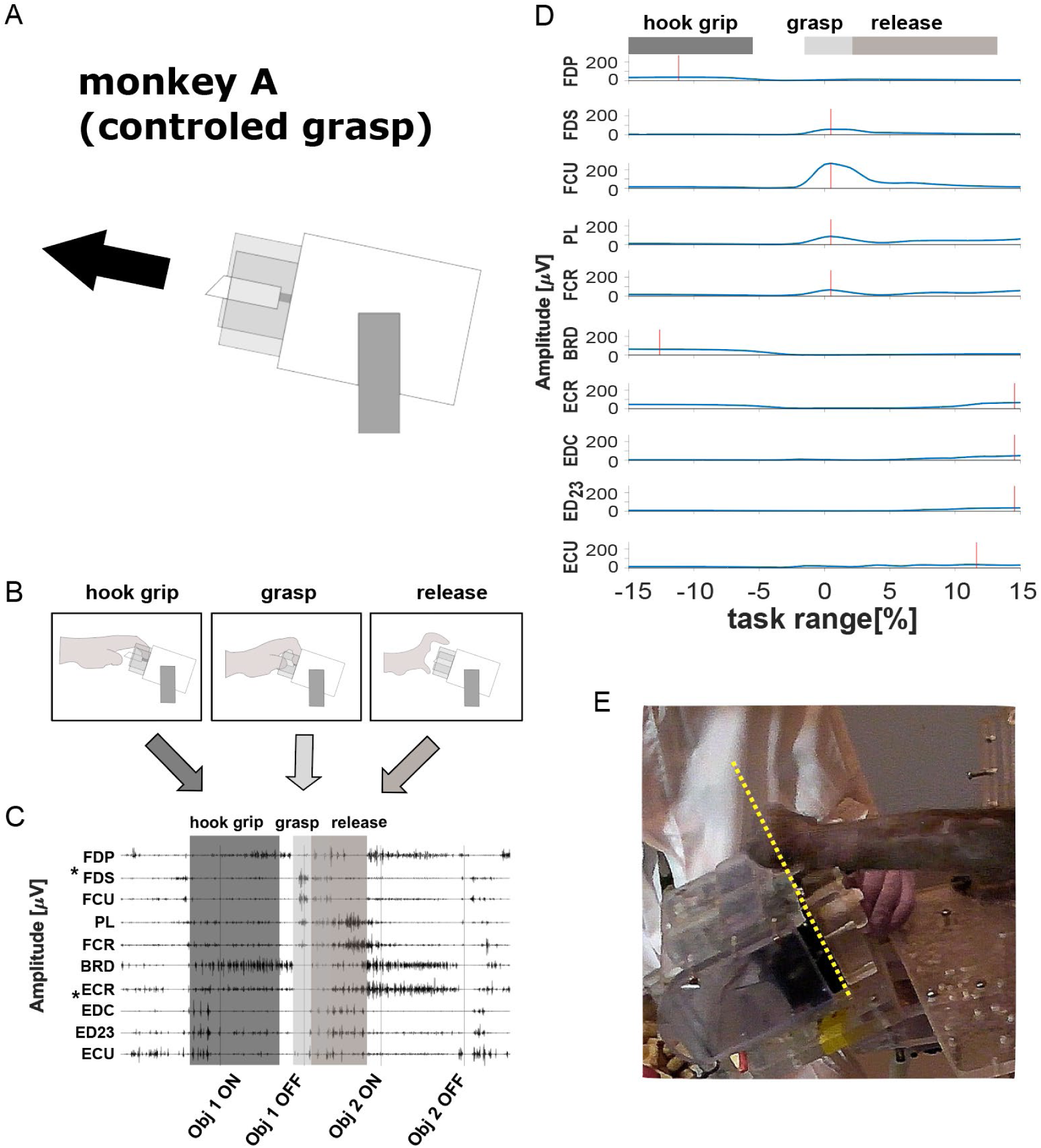
Experimental set-up, task sequence and typical EMG (monkey A). (A) Schematic of the task object using a rod requiring monkey A to perform a controlled grasp. (B) Schematic of the behavioral sequence (hook → grasp → release). (C) Typical EMG traces of a control session (high-pass filtered) for all recorded muscles. Gray boxes represent the task sequence. Obj 1 ON: start of the hold period of object 1. Obj 1 OFF: end of the hold period of object 1, i.e., object release. Obj 2 ON: start of the hold period of object 2. Obj 2 OFF: end of the hold period of object 2, i.e., object release. Tendons of the muscles marked with * were cross-transferred. (D) Rectified and smoothed EMG for all recorded muscles (average for one recording session; amplitude [μV] over task sequence [%]). Horizontal bars illustrate the corresponding behavioral periods; red vertical lines indicate peak amplitude for each muscle). (E) The time the monkey spent on the left side of the yellow dotted line while moving from object 1 to object 2 was measured and used to quantify the maladaptive behavior.

**Figure 4:**
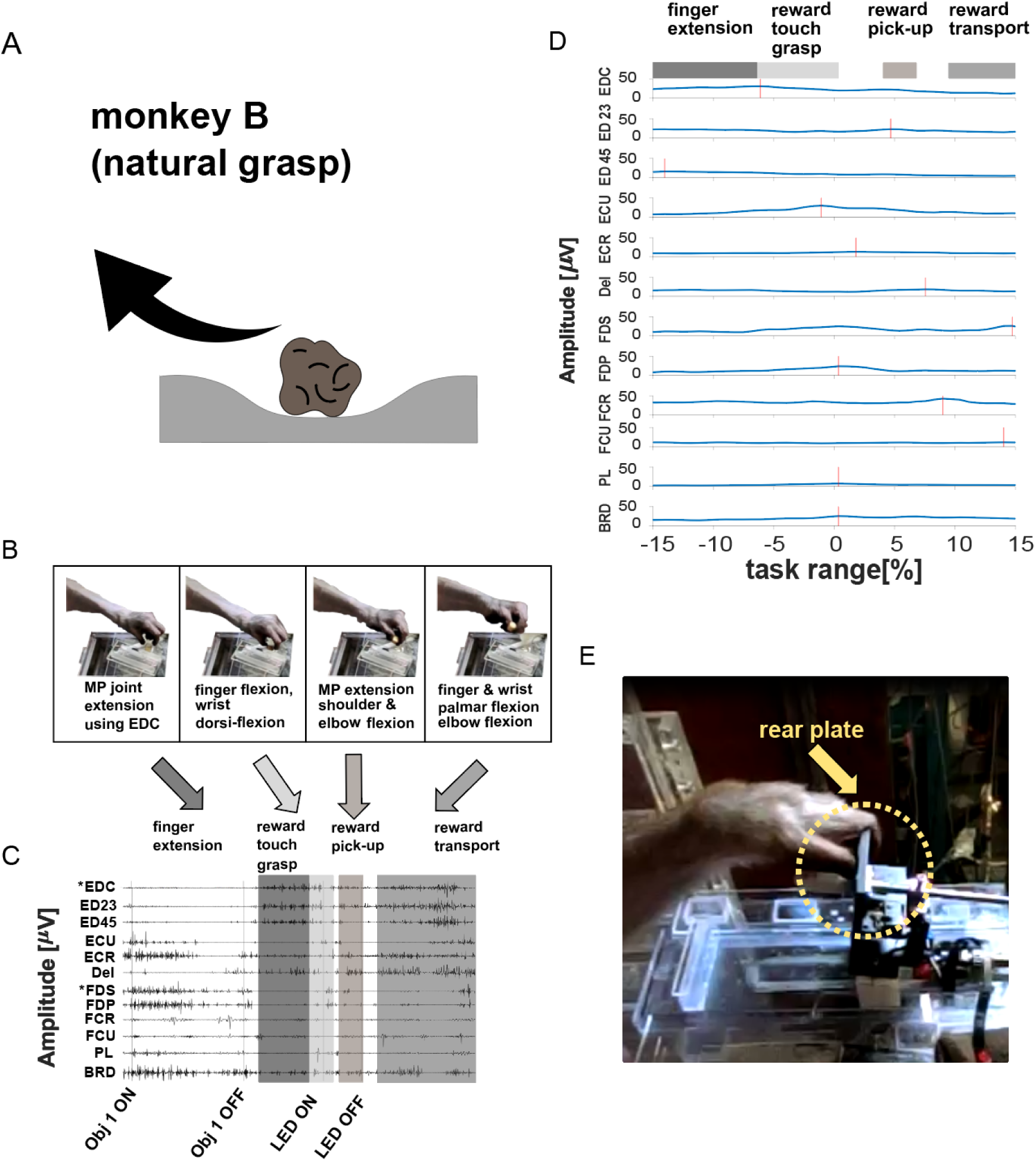
Experimental set-up, task sequence and typical EMG (monkey B). (A) Schematic of the task requiring monkey B to pick up food from a groove allowing for a more natural grasp. (B) Schematic of the task sequence (picking up food). (C) Typical EMG traces of a control session (high-pass filtered) for all recorded muscles. Gray boxes represent the task. Obj 1 ON: start of the hold period of object 1. Obj 1 OFF: end of object 1’s hold period, i.e. object release. LED ON: approximate start of the food touch. LED OFF: approximate time of food retrieval. Tendons of the muscles marked with * were cross-transferred. (D) rectified and smoothed EMG for all recorded muscles (average for one recording session; amplitude [μV] over task sequence [%]). Horizontal bars illustrate the corresponding behavioral periods, red vertical lines indicate peak amplitude for each muscle). (E) Example for maladaptive behavior in monkey B. The time the monkey spent in contact with or behind the object plate was measured and used to quantify the maladaptive behavior.

Once the monkeys had mastered their respective tasks, we recorded control sessions and then performed the crossed TT. Post-surgery, and once the monkeys had fully recovered and were able to perform the task independently (∼3–4 weeks), EMG recordings and behavior were resumed.

### Functional Recovery Follows a Biphasic Trajectory

Despite the now reversed roles of the transferred muscles, the monkeys were able to recover their grasping performance within two months, assessed by both the return of key behavioral metrics to pre-surgical levels (Fig. 5A, D) and their consistent ability to perform the task independently (Supplementary Videos S3, S4, S7). However, this recovery was not immediate. The monkeys spent a period of 1-2 weeks in their home cages for recovery, followed by several weeks of assisted task practice. Formal recordings began once the monkeys could perform the task consistently without assistance. The initial post-surgical period was defined by a phase of significant motor impairment, primarily characterized by off-target reaching trajectories (Fig. 5B, E) and prolonged grip formation times (Fig. 5A, D). These off-target reaching movements included inefficient, ‘explorative’ trajectories in Monkey A and target overshoots in Monkey B (Supplementary Videos S2, S3, S6). This recovery process was quantified across several behavioral and kinematic metrics (Fig. 5). A comparison of pre-TT behavioral variability on the grip formation task (Fig. 5D) revealed that Monkey B was substantially more variable (CV = 81.93%) than Monkey A (CV = 46.62%). This difference in variance was highly statistically significant (Ansari-Bradley test, p < 0.0001), confirming a difference in baseline stability between the subjects likely due to the less constrained nature of Monkey B’s task. Post-surgery, variability in grip formation time initially increased dramatically for Monkey A (Early Phase CV = 132.98%), while remaining high for Monkey B (Early Phase CV = 76.46%). In the late phase, Monkey A’s variability stabilized slightly above baseline (Late Phase CV = 54.79%), whereas Monkey B’s variability increased further, exceeding its pre-TT level (Late Phase CV = 96.59%), suggesting persistent inconsistency in grasp timing for this animal. Variability patterns differed for the other metrics: Monkey A’s pull time variability decreased below baseline in the mid and late phases (Pre CV = 64.97%, Mid CV = 41.30%, Late CV = 43.01%), indicating a refinement of this action, while Monkey B’s grasp aperture variability remained consistently low throughout recovery (Pre CV = 26.37%, Early CV = 23.80%, Mid CV = 19.78%, Late CV = 19.35%). The initial post-surgical period was defined by motor performance that we classify as maladaptive due to the persistence of inefficient, off-target reaching and prolonged movement durations. First, task-related grip formation times were significantly longer immediately post-TT (Monkey A: 197.7 ±92.2 vs. 660.6 ±221 ms, p = 0.014; Monkey B: 169.7 ±14 vs. 316.3 ±31.9 ms, p = 0.016), taking approximately 40 days for Monkey A and 48 days for Monkey B to return to and stabilize at pre-surgical levels (Fig. 5A, D). Second, the duration of off-target reaching was substantially elevated, stabilizing approximately after 70 days in Monkey A and 55 days in Monkey B (Fig. 5B, E). Following this phase, motor performance showed a gradual recovery in efficiency (Fig. 5C) and kinematics (Fig. 5F). Ultimately, overcoming these initial maladaptive behaviors through the gradual refinement of motor performance led to the stabilization of grasping by the end of the experimental period.

**Figure 5:**
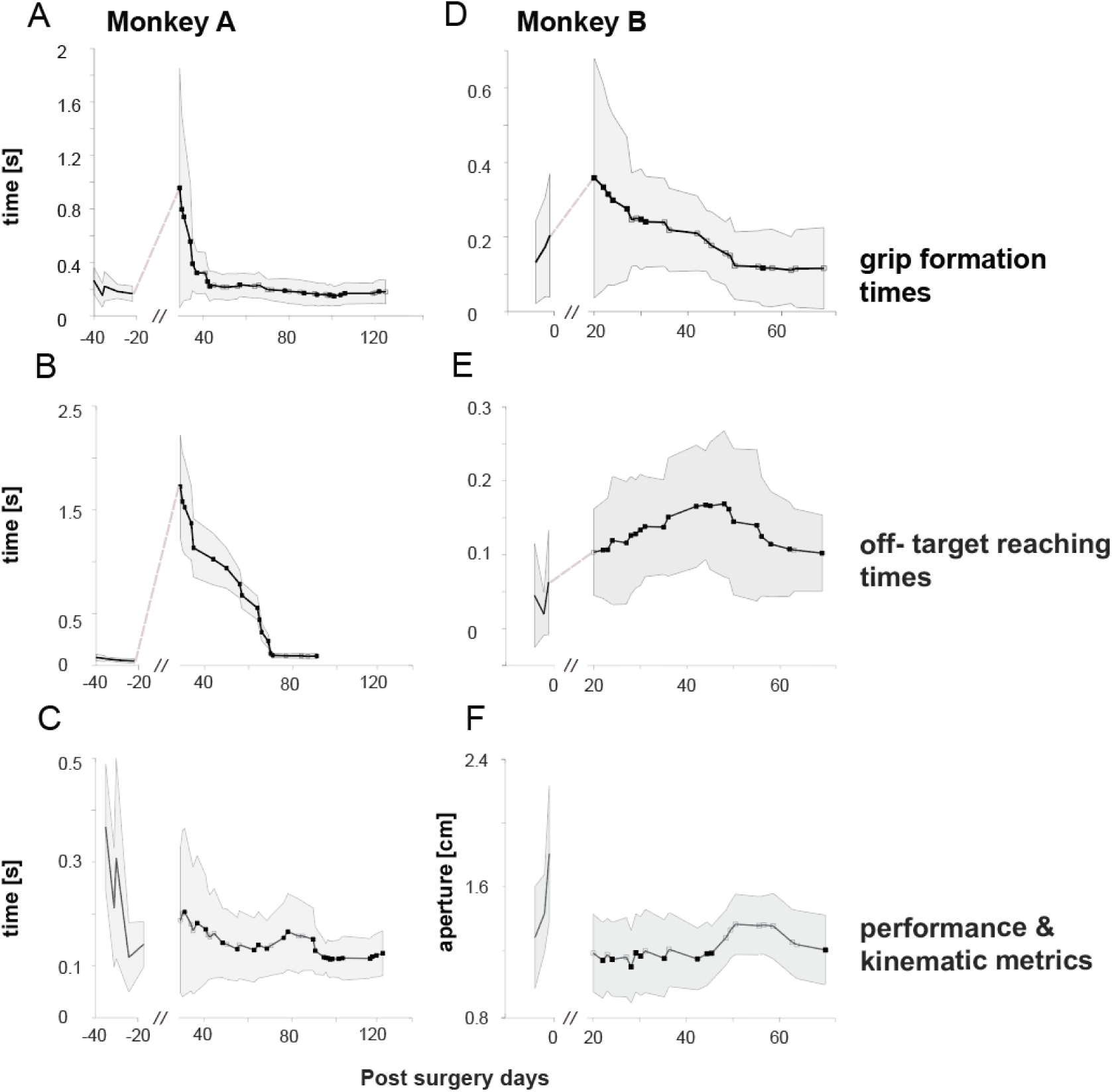
Behavioral and kinematic metrics of motor recovery. (A, D) Grip formation times (mean ± SD; n = 20 trials) for Monkey A (A) and Monkey B (D). (B, E) Duration of off-target reaching movements (mean ± SD; n = 10 trials) for Monkey A (B) and Monkey B (E). (C) Pull time duration for Monkey A. (F) Grasp aperture size for Monkey B. Filled squares indicate significant difference from pre-TT baseline (p < 0.05, two-sample t-test). All data are plotted over days relative to tendon transfer (TT).

### A two-phase adaptation is observed in the EMG activity of individual muscles

Initially, we investigated whether EMG activity exhibited any changes post-TT. Given that we interchanged the tendons of an antagonistic muscle pair, functionally rendering the FDS an extensor and the EDC a flexor (as confirmed mechanically, Fig. 2), a biomechanically sound adaptation would be to use the former finger extensor (EDC) for finger flexion, and the former finger flexor (FDS) for finger extension during a grasping task. We therefore analyzed EMG activity from both transferred and non-transferred muscles to determine if these expected functional changes occurred. Figure 6 shows a comparison of EMG activity profiles for the two transferred muscles, EDC and FDS, in Monkey A (Fig. 6A–D). Prior to surgery, the control showed distinct and contrasting EMG profiles in both muscles. For instance, peak activity in EDC (▾) occurred long after completion of the grasping action (indicated by vertical dashed lines at 0% task range) at 15% task time (Fig. 6A), coinciding with the animal pre-shaping its hand to prepare for the subsequent grasp. Whereas peak activity in FDS (▽) was observed immediately after completion of the grasping action (0.49%; Fig. 6C). The question we posed was whether the post-surgery activity of EDC resembled its original profile (without adaptation to surgery) or the activity of FDS (the anticipated profile following surgical adaptation), and vice versa. Our findings suggested that EMG activity patterns largely shifted in a manner consistent with the new mechanical function imposed by the transfer. Early after TT, peak activity of EDC occurred at −0.58% task time (Fig. 6B; days 29 [red line] and 64 [orange]), which aligns with peak activity of FDS prior to TT (Fig. 6C; black line). Also, for FDS, peak activity occurred at 11% task time (Fig. 6D; ▾), which aligned with peak activity of EDC (Fig. 6A).

**Figure 6:**
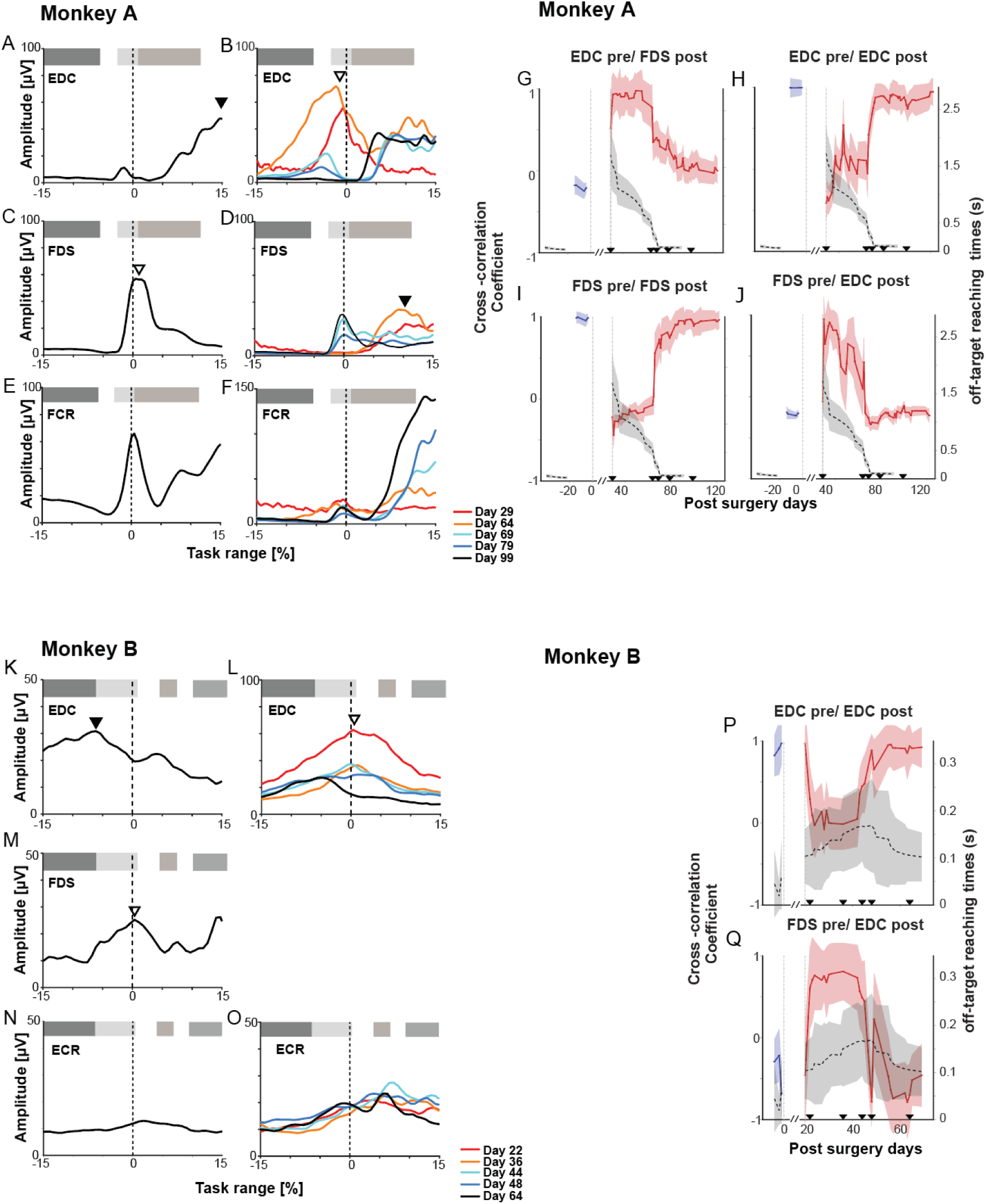
Temporal EMG profiles and cross-correlation analysis. Temporal EMG profiles for Monkey A (left, A-J) and Monkey B (right, K-Q). (A-F, K-O) Average EMG activity profiles aligned to task events (0%; object release for A, food touch for B). Shaded envelopes represent standard deviations. Triangles indicate peak activity during extension (▼) or flexion (▽). Colored traces denote post-surgery landmark days. (A-D, K-M) Comparison of transferred muscles. Note the temporal shift in the post-surgery profile (B, L) relative to the pre-TT baseline (dashed lines). (E-F, N-O) Profiles of non-transferred muscles. (G-J, P-Q) Zero-lag cross-correlation coefficients between post-surgery EMG profiles and the pre-TT baseline profiles plotted over time. (H, I, P) Correlation coefficients calculated against the muscle’s *own* pre-TT baseline. (G, J, Q) Correlation coefficients calculated against the *antagonist’s* pre-TT baseline (e.g., Post-EDC vs. Pre-FDS). Black dashed lines on the right y-axis indicate behavioral error metrics (Off-target reaching time for Monkey A; Contact duration for Monkey B; gray shading represents SD). The // represents the recovery period.

This was corroborated by the results in Monkey B (Fig. 6, panels K-M). Although the EMG patterns of both muscles varied significantly as the two monkeys performed different types of grasping actions, we found that the EMG of EDC peaked at −6.12% before (Fig. 6K; ▾) and 0.35% and 1.05% after TT (Fig. 6L; ▽, days 22 and 36, respectively), closely matching the timing for FDS control data (Fig. 6M). The corresponding analysis for FDS could not be performed in Monkey B, since the EMG signal of FDS was lost early after TT (see Methods for details). We were able to extend our recordings to 122- and 64-days post-TT for monkeys A and B, respectively. This allowed us to examine whether the adaptations observed in the early period were consistent throughout the experimental period. These additional days (Fig. 6B, D; 69 [cyan], 79 [blue], and 99 [black] post-TT; see Fig. S1 for recordings from all days) showed that the EMG activity profiles for both muscles unexpectedly returned to their pre-TT state. For example, as already described, the EMG signal of EDC at days 29 and 64 shared the characteristics of FDS, but by the following day (only 5 days later, day 69 [light blue] post-TT) already exhibited pre-TT characteristics of EDC, continuing as such until the end of the recording period (day 79 [blue] and 99 [black]). This was confirmed for the EDC muscle of Monkey B (Fig. 6K, L; Fig. S1U-AO for recordings from all days).

To relate these neural changes to functional recovery, we overlaid the behavioral error metric (off-target reaching time; gray dashed lines) onto the cross-correlation plots (Fig. 6G–J). Figure 6H (red line) illustrates cross-correlation coefficients between the EDC EMG profile post-TT (EDC-post) and the original EDC control data (EDC-pre) in Monkey A. Figure 6I shows coefficients for EDC-post and FDS-pre correlations in the same monkey. The coefficient corresponding to the original profile decreased to between −0.3 and 0.2 over approximately 65 days (Fig. 6H). In contrast, the coefficient corresponding to the expected EMG profile exhibited an inverse pattern, with low coefficients pre-TT and a high coefficient (of 0.9) post-TT (Fig. 6J). Cross-correlation analysis for FDS (Fig. 6G, I) revealed a similar pattern. Cross-correlation coefficients for FDS-pre- vs. FDS-post analysis declined from 1 to negative values, only to rebound to their original values within a month (day 79; Fig. 6G). This outcome aligns with our previous observations (Fig. 6B, D), and were corroborated by the results from Monkey B (Fig. 6K-Q). Here, cross-correlation coefficients dropped from +0.95 to +0.1 for EDC-pre vs. EDC-post comparison, and increased from −0.5 to 0.9 for the FDS-pre vs. EDC-post comparison (Fig. 6P, Q). After 42 days post-TT, cross-correlation coefficients started to gradually increase or decrease to +0.9 and −0.7, respectively. This indicates that the anticipated initial changes in EMG activity profiles are not permanent. Instead, they revert to their original patterns two months post-TT. A supplementary analysis of the time lag at peak correlation (Supplementary Figure S1) confirmed that the optimal lag for individual muscles also fluctuated significantly during the early adaptive phase before stabilizing, reflecting the process of temporal re-coordination of individual muscle commands.

Interestingly, changes in EMG pattern were not isolated to the transferred muscles; non-transferred muscles exhibited a variety of complex adaptations (Figs. 6, S2-S4). A comprehensive overview of the average EMG profiles for all recorded muscles across all recording sessions and landmark days is provided in Supplementary Figures S2 (Monkey A) and S3 (Monkey B), respectively. As shown in these figures, many agonists adapted in concert with their transferred synergist, following the same two-phase “swap-and-revert” pattern seen in Figure 6. In Monkey A, these included extensors ED23 and ECU, while in Monkey B, they included ED23, ED45, and ECU (see Fig. S2 for details). In contrast, other muscles showed patterns that were incompatible with a simple swap. For example, the non-transferred flexor carpi radialis (FCR) in Monkey A showed a distinct adaptive profile characterized by a drastic initial decrease in one activity peak and a consistent increase in a later peak (Fig. 6E, F). A similar incompatible pattern was seen in palmaris longus (PL) (Fig. S1H, I, R, S). Finally, some muscles, like the extensor carpi radialis (ECR) in Monkey B, remained relatively stable post-surgery (Fig. 6N, O). These diverse patterns in non-transferred muscles do not necessarily contradict a modular control strategy (d’Avella et al., 2003; Bizzi and Cheung, 2013); rather, they likely reflect the fact that different muscles are primary members of different synergies (i.e., they possess the highest weightings within specific modules; e.g., FCR and PL in Synergy C), each undergoing its own task-specific adaptation.

### Adaptation occurs through modulating the activation of stable muscle synergies

To determine whether the CNS adapted by fractionating muscle control or modulating existing modules, we extracted muscle synergies using non-negative matrix factorization (NMF) (Lee and Seung, 1999). Four synergies accounted for >80% (Cheung et al., 2012) of the variance in both monkeys (see Supplementary Figure S5 for VAF plots; >85% for both monkeys; the results of the original and shuffled data sets are shown. See also the synergy weights [*W*] and activation profiles [*C*] for both monkeys and all synergies in figs. S6 and S7, respectively). We first focused on the two primary synergies responsible for the main task axis: the finger flexor synergy (Synergy A) and the finger extensor synergy (Synergy B) (Fig. 7).

**Figure 7:**
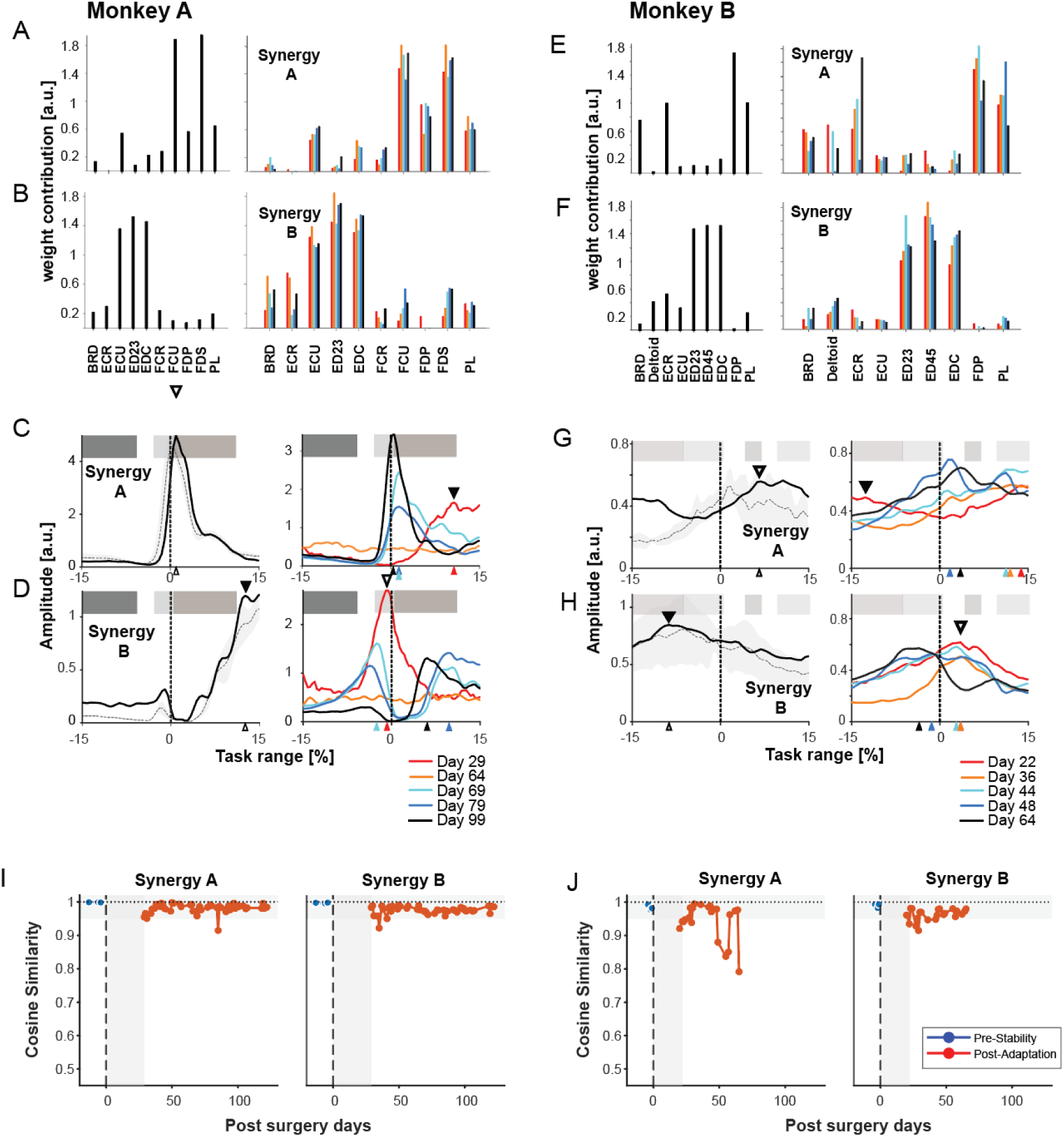
Spatial structure and temporal activation of Primary Synergies. Analysis of the two primary synergies: Synergy A (Flexor) and Synergy B (Extensor). (A, B, E, F) Spatial synergy weights (*W*) showing the contribution of each muscle to the synergy. Bar plots represent the average weights across all recording days. (C, D, G, H) Temporal activation profiles (*C*) aligned to task events (0%). Dashed lines with shaded tubes indicate the average pre-TT EMG profiles of the key contributing muscles (FDS, EDC, FDP) for visual comparison with the synergy profile. Symbols and alignment are as described in Figure 6. (I, J) Quantification of spatial stability. Cosine similarity of spatial synergy weights (*W*) calculated between individual recording days and the pre-TT average. Blue markers indicate pre-TT control days; Red markers indicate post-TT days. The horizontal gray shaded region (0.95-1.0) denotes the range of high baseline stability.

We first analyzed whether the spatial structure of these primary synergies changed. To distinguish genuine adaptation from natural variability, we established a baseline by calculating the cosine similarity of spatial synergy weights (*W*) across all pre-surgery days (Fig. 7I, J, blue traces). This revealed remarkably high stability (>0.99). Post-surgery, the core structure remained conserved. The cosine distance between pre- and post-surgery weights for the same synergy pairs was low (Monkey A: 0.03 ± 0.03; Monkey B: 0.09 ± 0.11), whereas for different pairs it was high (>0.60). This confirms that the CNS maintained the stable, neurally constrained building blocks of the primary flexor and extensor modules despite the altered biomechanics.

In contrast to the stable spatial structure, the temporal activation profiles of these primary synergies underwent drastic changes. In Monkey A, the flexor Synergy A (originally peaking at +0.97% task time) and extensor Synergy B (originally peaking at 12.62%) swapped their timing profiles shortly after surgery (Fig. 7C, D). Specifically, the extensor synergy shifted to peak during the flexion phase, and vice versa. This “swap” persisted during the early impairment phase before reverting toward the original timing in the late phase. A similar “swap-and-revert” pattern was observed in Monkey B (Fig. 7G, H), where the extensor Synergy B shifted from a pre-surgery peak at −8.75% to a post-surgery peak of +3.5% (matching the flexor phase) before recovering.

To formally quantify the quality of this reversion, we compared the pre-surgery activation profiles with those from the final day of recording. We found that while the temporal shape was highly preserved (Cosine Similarity >0.90), the profiles remained statistically distinct (Permutation Test, p < 0.0001; Fig. S8). This confirms that the recovered motor program represents a “good enough” functional approximation rather than a perfect mathematical restoration of the baseline.

We next examined the secondary synergies involved in wrist control and stabilization (Fig. 8). In contrast to the coherent “swap” seen in the primary synergies, these modules exhibited distinct, task-specific adaptations.

**Figure 8:**
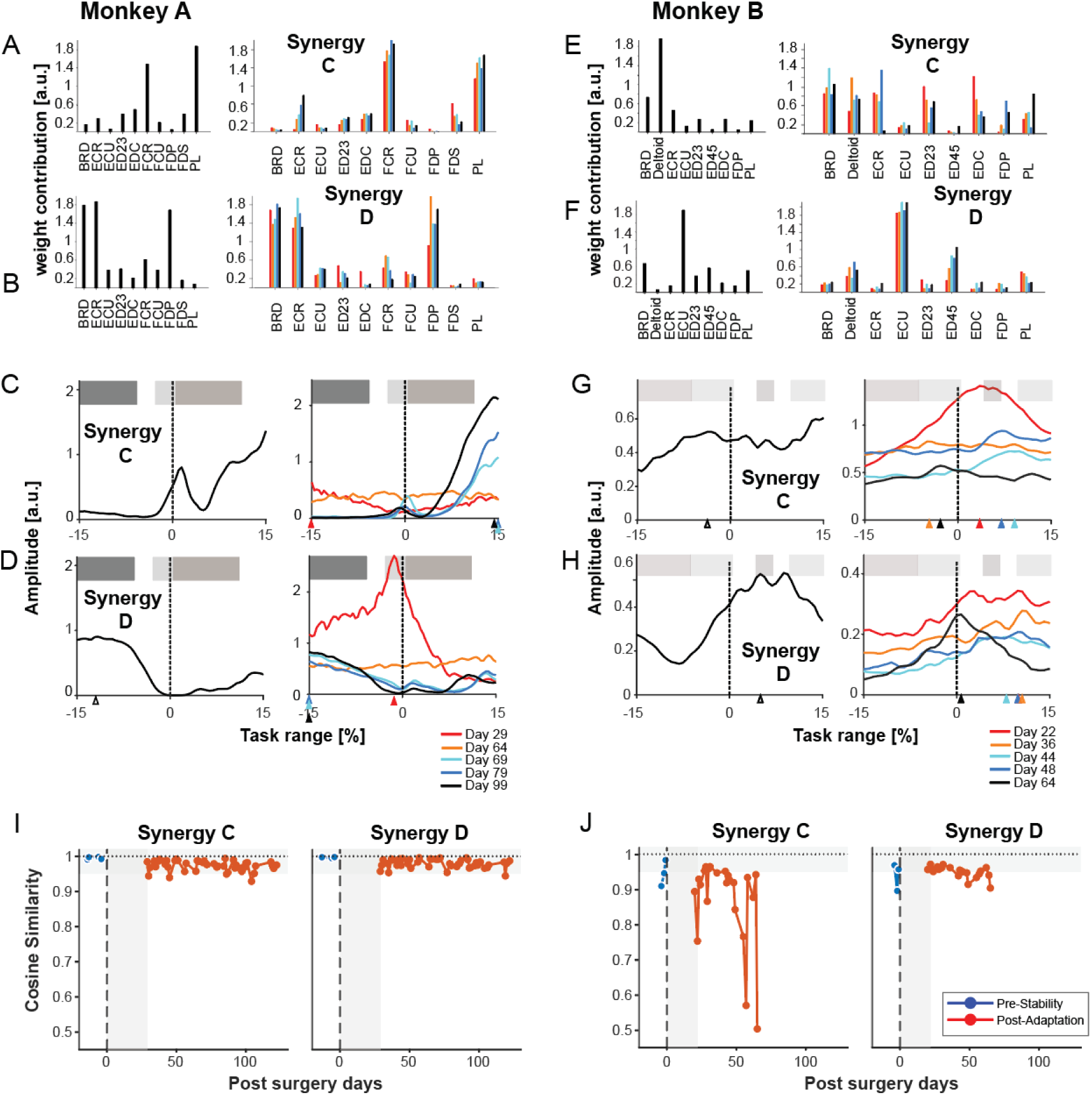
Analysis of Secondary Muscle Synergies. (A-H) Analysis of Secondary Synergies C (Wrist Flexor) and D (Wrist Extensor). (A, B, E, F) Spatial synergy weights (*W*) showing the contribution of each muscle. (C, D, G, H) Temporal activation profiles (*C*) aligned to task events (0%). Colored traces denote post-surgery landmark days. Layout and symbols are as described in Figure 6. (I, J) Quantification of spatial stability for Synergies C and D. Cosine similarity of spatial weights calculated between individual recording days and the pre-TT average. Blue markers indicate pre-TT control days; Red markers indicate post-TT days. The horizontal gray shaded region (0.95-1.0) denotes the range of high baseline stability.

In Monkey A, Synergy C (a wrist flexor synergy involving PL and FCR) showed a temporal shift that was incompatible with a simple swap (Fig. 8C). While its spatial weights remained stable (Fig. 8A, I), its activation peak shifted from the post-grasp phase (+1.46%) to the pre-grasp phase, increasing in magnitude over time. This pattern corresponds to the recruitment of wrist flexors to enable the compensatory tenodesis grasp. Synergy D (wrist extensor) underwent a single notable change on day 29 but quickly reverted (Fig. 8D). In Monkey B, Synergy C showed no discernible trend (Fig. 8G), while Synergy D exhibited a complex pattern of immediate change followed by a gradual shift in the late adaptation period (Fig. 8H). Notably, the activation amplitude of Synergy D remained significantly suppressed compared to baseline (Fig. 8H; Fig. S6H). This suppression of the wrist extensor likely reflects the specific biomechanical requirements of the tenodesis strategy employed by Monkey B: relaxing the wrist extensors facilitates the necessary wrist flexion for hand opening. These independent changes in secondary synergies likely served to reinforce the overall functional recovery driven by the primary modules.

Again, the characteristics of these changes in activation profiles of muscle synergies were also quantitatively confirmed by cross-correlation analysis of all four muscle synergies in both monkeys. The coefficients were plotted over the course of experimental days in relation to tendon surgery together with the overlaid behavioral error metric (off-target reaching time; gray dashed lines) (Fig. 9). In this analysis, the activation profile of each synergy was cross-correlated with either the one from synergy A (finger flexor synergy; Fig. 9C, D, G, H) or synergy B (extensor synergy; Fig. 9A, B, E, F) before TT. As previously demonstrated for the FDS and EDC muscles, the temporal activation profiles of muscle synergies A and B (flexor and extensor) displayed a distinct pattern. After cross-correlation with their own control data, both the flexor and extensor synergies showed coefficients of 1 prior to surgery. These became negative for 66 days post-surgery before reverting close to their original values (Fig. 9B, C). However, after cross-correlation with control data of the antagonistic muscle synergy (i.e., extensor for flexor synergy and vice versa, Fig. 9A, D), the pattern was reversed. Coefficients started with negative values prior to TT and shifted to positive coefficients near 1 shortly after surgery. In Monkey A, coefficients returned to negative values around day 66 post-TT. In Monkey B, this reversal occurred earlier, around day 48. In short, these results mirror our findings between the transferred muscles (Fig. 6G, J). Furthermore, to determine if low correlations were driven by simple timing mismatches, we analyzed the optimal time lag yielding the maximum cross-correlation (Supplementary Figure S9). A positive lag indicates the post-surgery activity is delayed relative to baseline, while a negative lag indicates it is advanced. This analysis revealed significant fluctuations in timing (switching between delays and advances) during the early and mid-adaptation phases, particularly around the ‘switch-back’ period, before stabilizing closer to zero lag in the late phase. For synergies C (main contributing muscles: ECR, BRD, DEL) and D (main contributing muscles: ECU), the changes in cross-correlation coefficients over time were markedly different. For synergy C, coefficients begin to steadily increase when cross-correlated with control data of the extensor synergy (synergy B, Fig. 10E), and decrease with the flexor synergy (synergy A, Fig. 10G). This suggests an increasing contribution to finger extension. In contrast, synergy D does not exhibit any specific pattern following TT.

**Figure 9:**
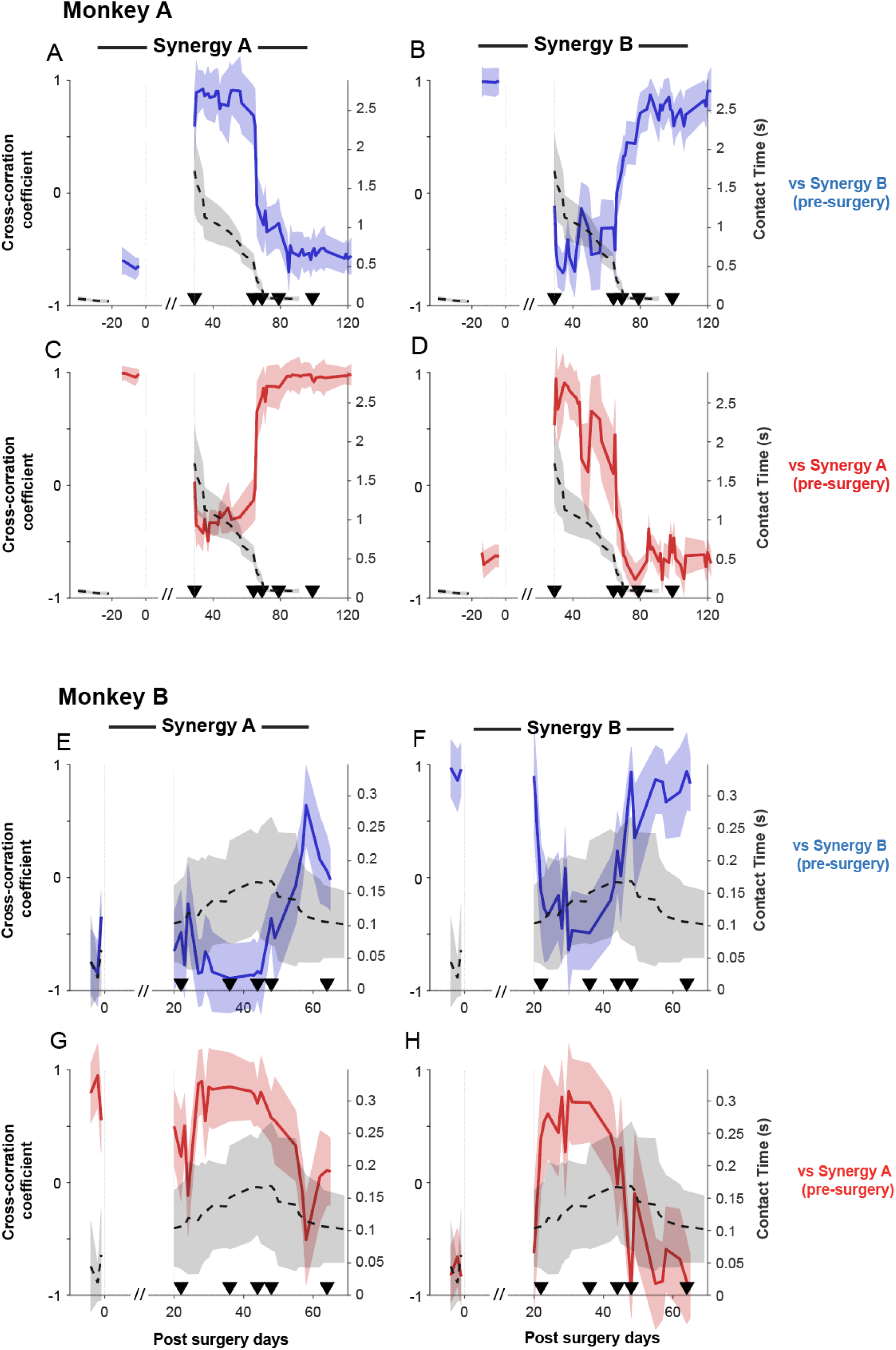
Cross-correlation analysis of Primary Synergy activation. Zero-lag cross-correlation coefficients plotted over post-surgery days for Monkey A (A-D) and Monkey B (E-H). Activation patterns of the Primary Flexor (Synergy A) and Primary Extensor (Synergy B) were cross-correlated with pre-TT baseline profiles. (Top Row: A, B, E, F) Correlations calculated against the pre-TT *Extensor* Synergy (Synergy B, blue traces). (Bottom Row: C, D, G, H) Correlations calculated against the pre-TT *Flexor* Synergy (Synergy A, red traces). (C, G) Correlation of Synergy A with its own pre-TT baseline. (B, F) Correlation of Synergy B with its own pre-TT baseline. (A, E, D, H) Cross-correlations between antagonistic synergies (e.g., A is Synergy A vs. Pre-Synergy B). Black dashed lines on the right y-axis indicate behavioral error metrics (gray shading represents SD). The // represents the recovery period. Triangles indicate landmark days.

**Figure 10:**
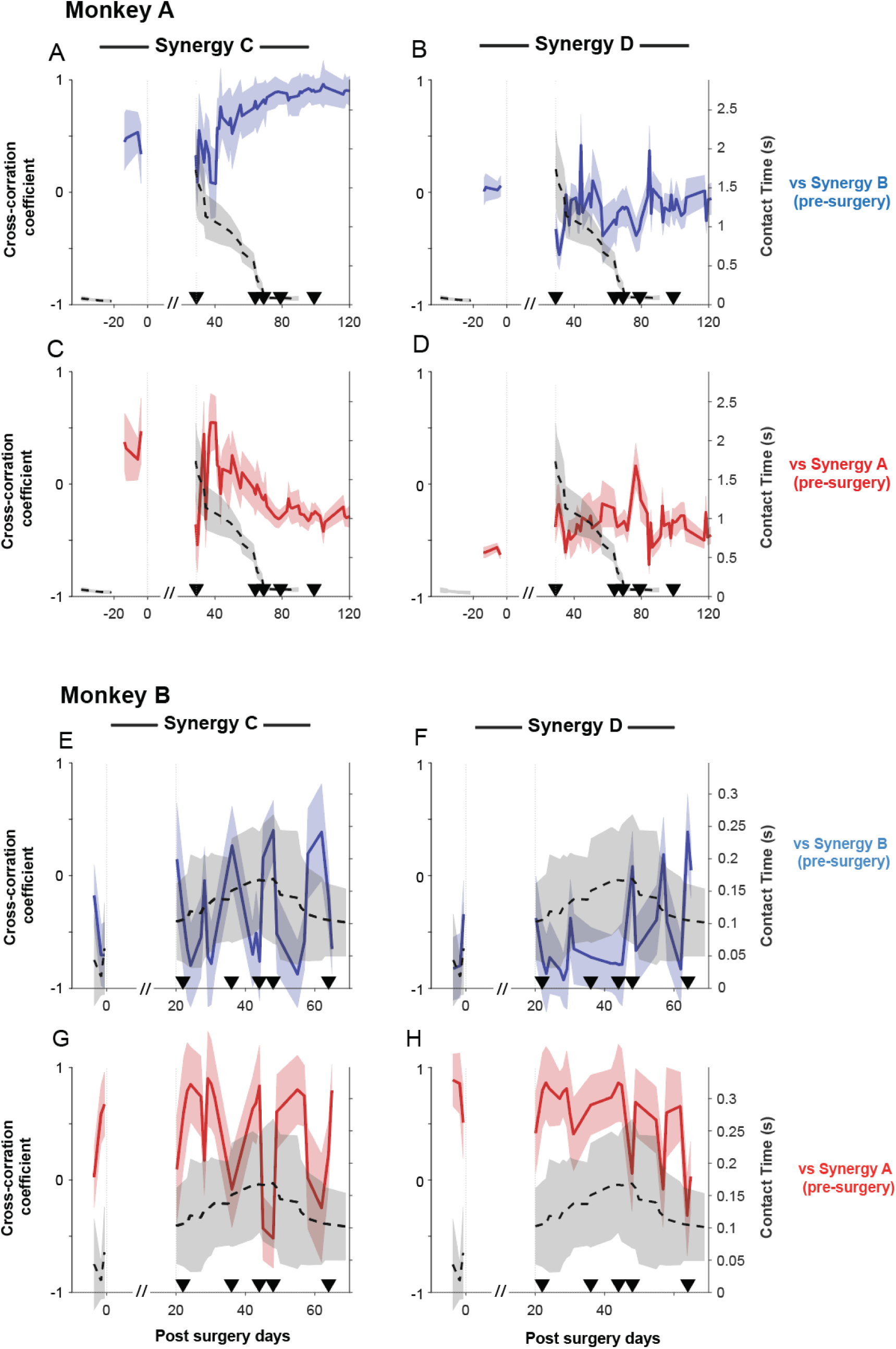
Cross-correlation analysis of Secondary Synergy activation. Zero-lag cross-correlation coefficients for the secondary synergies (C and D) plotted over post-surgery days for Monkey A (A-D) and Monkey B (E-H). Activation patterns were cross-correlated with the pre-TT profiles of the Primary Synergies to assess changing affiliations. (Top Row: A, B, E, F) Correlations calculated against the pre-TT *Extensor* Synergy (Synergy B, blue traces). (Bottom Row: C, D, G, H) Correlations calculated against the pre-TT *Flexor* Synergy (Synergy A, red traces). (Left Columns: A, C, E, G) Synergy C correlations. (Right Columns: B, D, F, H) Synergy D correlations. Black dashed lines on the right y-axis indicate behavioral error metrics.

In Monkey B, this cross-correlation analysis revealed a more varied, or differential, pattern of adaptation across the four synergies (Figs. 9 and 10). The primary extensor, Synergy B, mirrored the results in Monkey A, showing a clear swap-and-revert pattern (Fig. 9F, H). In contrast, Synergy A did not show a clear reversal; its correlation coefficients gradually converged towards zero, likely due to the absence of the FDS EMG signal in the analysis (Fig. 9E, G). The secondary synergies, Synergy C and D, showed no discernible trend (Fig. 10E-H).

In summary, both monkeys exhibited a distinct two-phase adaptation following TT. We define the ‘early phase’ as the period from initial post-surgical recovery up to the reversal of the swapped activation patterns (approximately days 20/29 to ∼65 post-TT, see Fig. 9), characterized by the transferred muscles/synergies adopting antagonistic temporal profiles. We also identify a ‘mid-adaptation’ phase covering the transitional period around the ‘switch-back’ event, characterized by high variability. The ‘late phase’ encompasses the period following this switch-back (after ∼day 66 post-TT), where original activation timings were largely restored. While this ‘reversion’ towards original activation timings in the late phase marked the abandonment of the initial maladaptive strategy, behavioral recovery involved more than just restoring original patterns. Concurrent with this neural shift in the primary synergies, a distinct compensatory strategy began to emerge, utilizing secondary synergies and biomechanical coupling to achieve functional hand opening, as detailed below.

### The Early Adaptation Phase is Characterized by a Maladaptive Neural and Behavioral Profile

The early adaptation phase was defined by a distinct neural strategy that correlated with significant behavioral deficits. First, the period dominated by off-target reaching movements and prolonged grip times (Fig. 5B, E) was precisely when the swapped activation of Synergy A and B was most prominent (Fig. 9; see behavioral overlay). This temporal link provides strong evidence that this initial ‘swap’ strategy was, in fact, maladaptive, as the flawed neural control directly underpinned the impairments in movement efficiency and precision.

Second, we found that the net activity of muscles representing certain muscle synergies (aggregated average EMG; aaEMG) showed distinct, synergy-specific changes over time (Fig. 11). A key feature was the evolution of the activity of finger extensor synergy (Synergy B), which appeared to undergo a neuromuscular ‘arms race’, a rapid escalation of antagonistic co-activation, in both animals, albeit on different timescales (Fig. 11B, J). In Monkey A, which struggled for a longer period to regain behavioral proficiency according to our subjective evaluation, our recordings now captured this process unfolding: we observed a steady and significant increase in Synergy B’s aaEMG post-TT until day 64 (p < 0.0001) before the strategy changed (Fig. 11B, F). In contrast, Monkey B, which adapted its behavior more rapidly according to our subjective evaluation, showed a different profile; its Synergy B activity peaked with a significant surge early in the recording period (day 22: p < 0.0001) and subsequently declined toward and below baseline (day 64: p < 0.001) (Fig. 11J, N). This suggests the initial, rapid escalation of the ‘arms race’ in this animal had already occurred and peaked prior to our first post-operative recording session. Therefore, the apparent discrepancy in the evolution of Synergy B is likely not a fundamental difference in strategy, but rather a reflection of the different observational windows and overall adaptation rates of the two animals.

**Figure 11:**
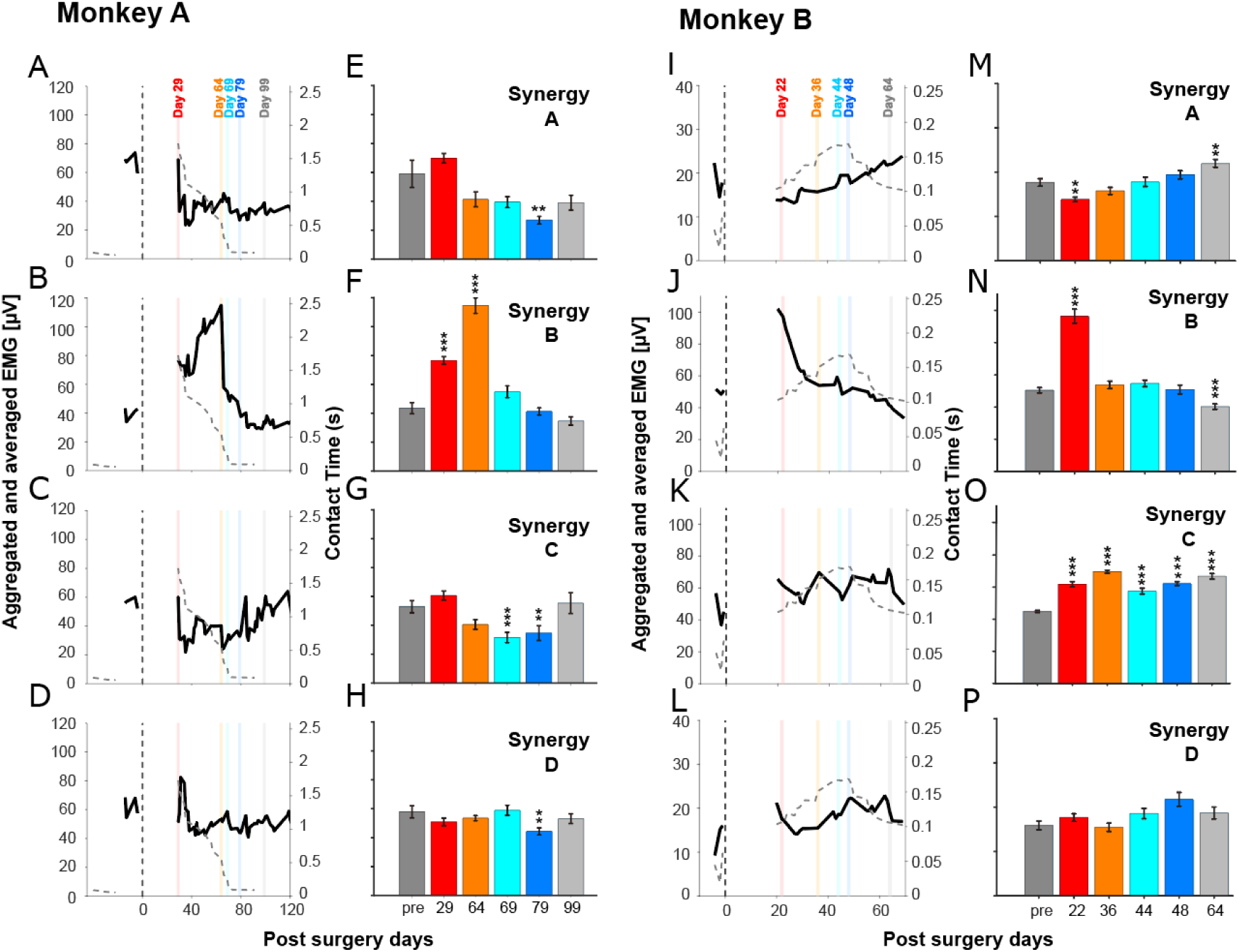
Aggregated and averaged EMG (aaEMG). Aggregated and averaged electromyography (EMG) activities for the main contributing muscles of each synergy for Monkey A (A-H) and Monkey B (I-P). (A-D, I-L) Time course of aaEMG activity (summed within ±15% task range) plotted over post-surgery days. Black dashed lines on the right y-axis indicate behavioral error metrics. (E-H, M-P) Bar plots showing the mean (± SEM) aaEMG for the pre-TT period (“pre”) and the five selected landmark days. Vertical colored bars on the time-series plots indicate the corresponding landmark days. Asterisks indicate significant difference from the pre-TT control period (*p < 0.01, **p < 0.001, ***p < 0.0001; two-sample t-test with Bonferroni correction).

For the finger flexor synergy (Synergy A), Monkey A showed a general decrease in activity (p = 0.0004 at day 79; Fig. 11A, E), while Monkey B showed a consistent and significant increase throughout the experiment (day 22: p = 0.0004; day 64: p = 0.0008). This was expected, as Monkey B’s Synergy A relied on FDP and PL to compensate for the loss of FDS function (Fig. 11I, M). Synergy C exhibited a general increase in activity during the post-TT phase in both monkeys (Fig. 11G, O), while Synergy D showed no significant differences in Monkey B and only minor changes in Monkey A (Fig. 11H, P). All statistical comparisons were made against the pre-TT control period using a two-sample t-test with Bonferroni correction (α = 0.01).

Taken together, these distinct patterns of aggregated EMG activity, especially the escalating co-activation within the conflicted Synergy B, a known marker of inefficient muscle recruitment (Thoroughman and Shadmehr, 1999; Osu et al., 2002), further illustrate that this early adaptive phase, which coincided with poor behavioral performance (Fig. 5), was characterized by an unstable and inefficient neural control strategy.

### Distinct Neural Implementations of a Compensatory Tenodesis Strategy

The final phase of adaptation was characterized by the development of a compensatory tenodesis strategy. This effect describes the passive coupling of finger and wrist movements due to the routing of tendons across multiple joints (Zajac, 1992): because the finger extensor tendons pass over the back of the wrist, actively flexing the wrist passively tightens these tendons, which in turn pulls the fingers into extension (Valero-Cuevas and Hentz, 2002; Cash and Jones, 2011). For Monkey B, we found direct kinematic evidence for the acquisition of this strategy. Post-surgery, the monkey learned to significantly increase finger extension at the metacarpophalangeal (MCP) joint (Fig. 12A) by concurrently flexing the wrist (Fig. 12B; p < 0.0001, ANOVA; α = 0.01). This coordinated movement pattern is characteristic of an active tenodesis effect. Furthermore, kinematic analysis revealed that this was a gradually learned skill rather than an immediate mechanical consequence. The trial-by-trial coupling between wrist angle and MCP angle (Fig. 13) was initially absent (R^2^ = 0.00) but strengthened over weeks, peaking in the mid-adaptation phase (R^2^ = 0.58). This period coincided with the stabilization of grasp aperture (Fig. 5F) and the resolution of the maladaptive neural patterns (Fig. 9). This kinematic strategy was supported by a precise neural implementation. Unlike the scaling strategy observed in Monkey A (see below), Monkey B did not rely on a massive increase in total muscle activity; the aggregated activation of its primary flexor synergy (Synergy A) showed consistent but moderate increases (Fig. 11I, M). Instead, the strategy was one of temporal refinement. The activation profile of Synergy A shifted to increase specifically during the pre-contact phase (Fig. 7G), providing the necessary wrist flexion for the tenodesis grasp. This was achieved by a precise sequential activation of muscles within the synergy, with the wrist flexor component (PL) peaking just before contact and the finger flexor component (FDP) peaking just after (Fig. S2). For Monkey A, which successfully restored its primary extensor synergy, the tenodesis grasp likely served as a similar compensatory driver. While we could not perform a kinematic analysis for this animal due to low-resolution video images, strong evidence for this strategy is provided by the neural data. We observed a clear temporal shift in the activation of its dedicated wrist flexor synergy (Synergy C). The peak of this synergy’s activation moved from occurring just after object contact to just before it (Fig. 8C), a re-timing well-suited to enable a tenodesis grasp. This increasing contribution is further supported by cross-correlation analysis, which shows its activation pattern became progressively more similar to that of the pre-TT extensor synergy over time (Fig. 10A; p < 0.05, two-sample t-test).

**Figure 12:**
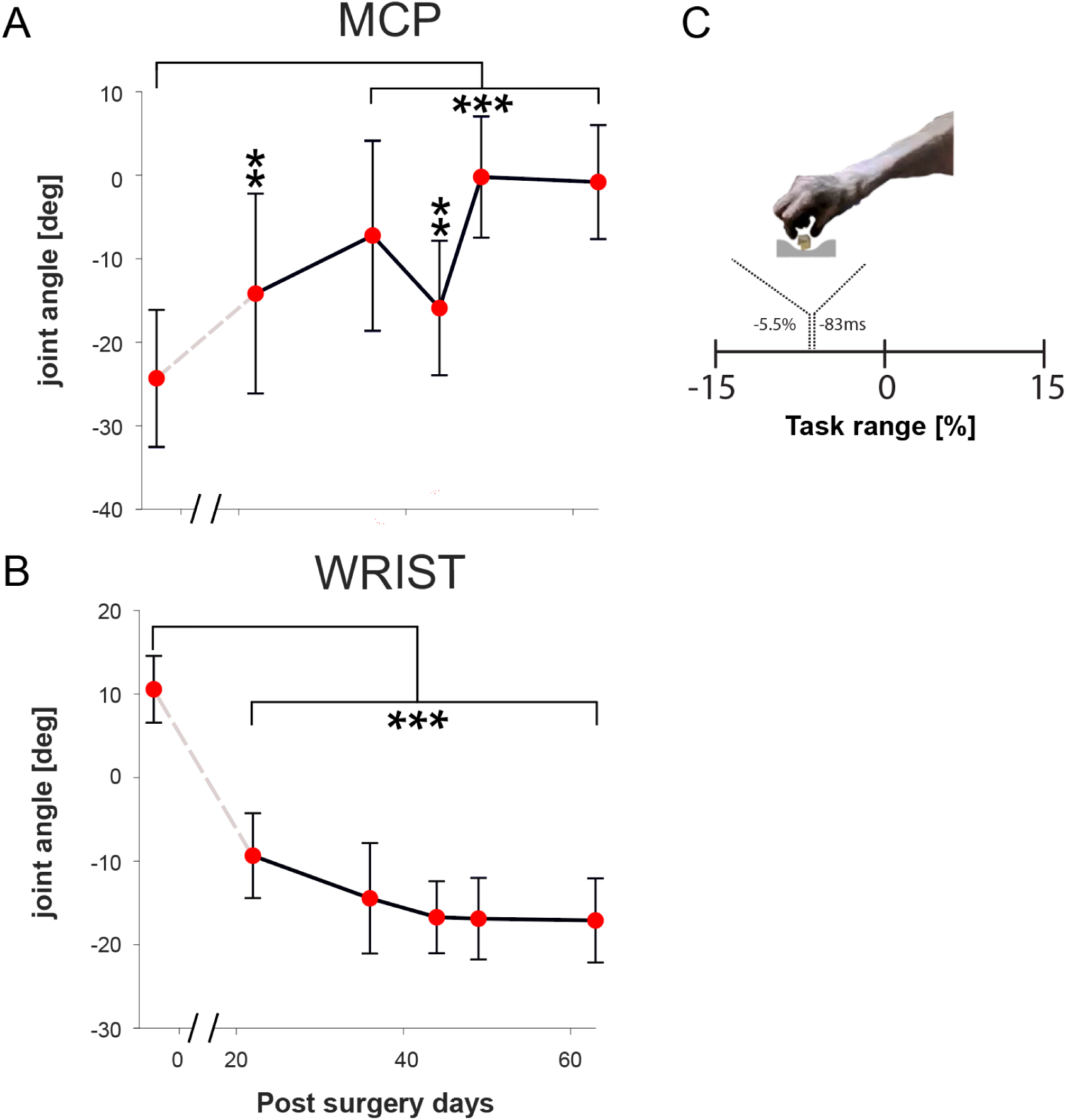
Kinematic analysis of joint angles (Monkey B). Changes in joint angles (mean of 20 trials ± SD) for each landmark day. (A) Metacarpophalangeal (MCP) joint angle. (B) Wrist joint angle. Asterisks indicate significant difference from pre-TT baseline (**p < 0.001, ***p < 0.0001; ANOVA). (C) Schematic indicating the timing of the kinematic snapshot relative to the task timeline (dotted line; 83 ms before food touch), capturing the hand configuration during the pre-shaping phase.

**Figure 13:**
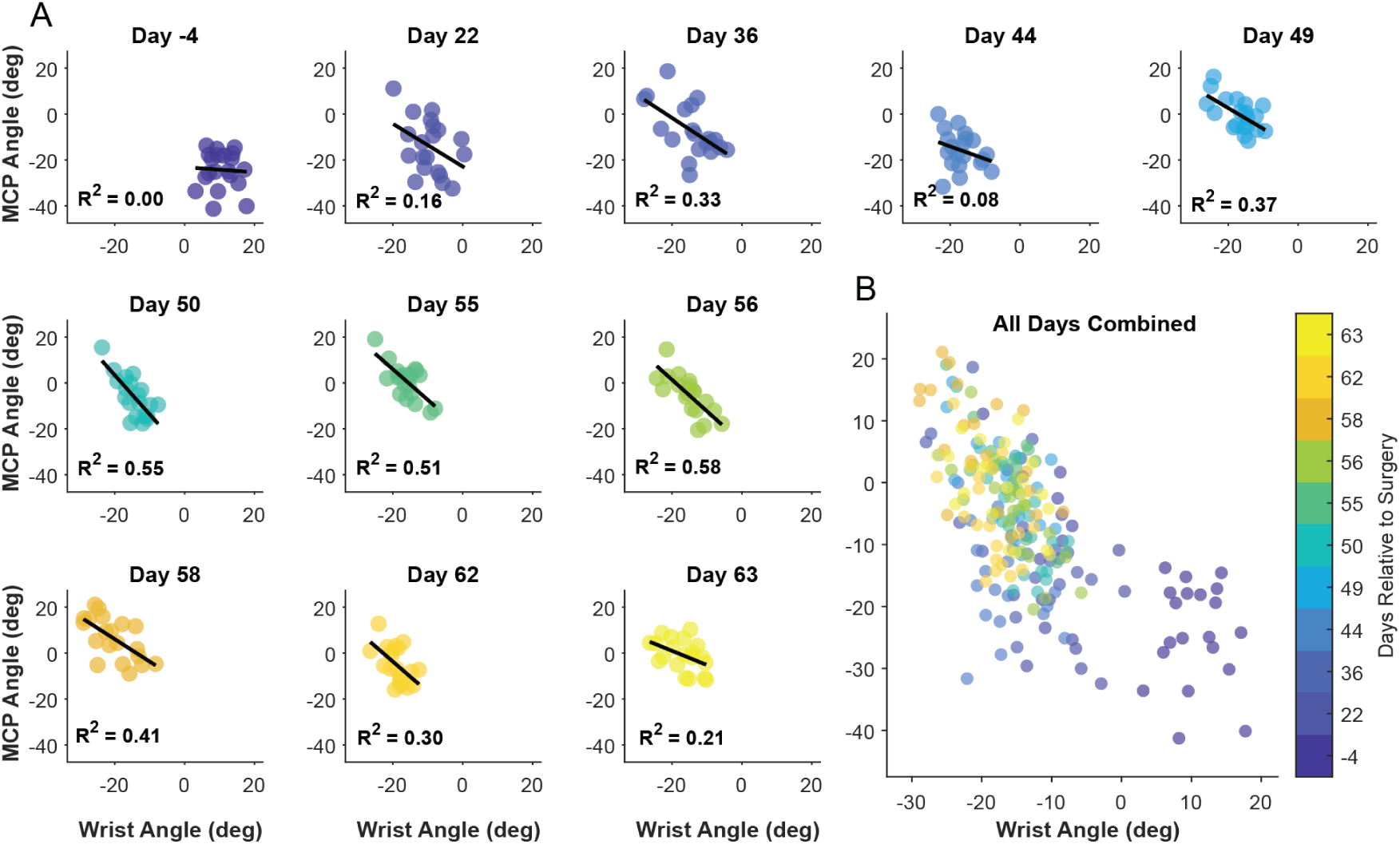
Kinematic analysis reveals gradual refinement of the compensatory tenodesis strategy over time in Monkey. **B.** (A) Each subplot shows the trial-by-trial relationship between wrist angle (X-axis) and MCP angle (Y-axis) for a single recording day (n=20 trials per day). Points are color-coded based on the day relative to surgery (colorbar). Pre-TT (Day -4), no correlation exists (R²=0.00). Post-surgery, a negative correlation emerges and strengthens over time, peaking around Day 56 (R²=0.58), indicating the learned exploitation of the tenodesis effect where wrist flexion predicts finger extension. The tightening of the scatter plots and increase in R² over weeks provide direct evidence for a gradual motor skill learning process. (B) All data points combined.

## Discussion

The central nervous system’s (CNS) response to profound musculoskeletal change is a fundamental problem in motor control. This study sought to determine whether the primate CNS adapts to such change by flexibly modulating stable muscle synergies or by developing more fractionated, independent muscle control. We found that the CNS initially defaulted to a modular strategy, repurposing entire synergies by swapping their activation timings. This simple solution, however, proved to be maladaptive, creating a mechanical conflict that impaired motor function under tendon transfer surgery used in this study. This early maladaptive phase was ultimately resolved through the gradual development of compensatory movements, leading to a “good enough” functional recovery. This multi-stage process, operating on different timescales, highlights the intricate balance between modularity and flexibility in neural adaptation.

### Neural Learning, Not Physical Healing, as the Rate-Limiting Factor

A crucial consideration in interpreting our long-term results is the potential confound of the tendon’s physical healing process. However, the surgical technique employed, a multi-weave Pulvertaft transfer, is designed to provide immediate mechanical strength comparable to that of a native tendon (Graham et al., 2023). While fibrous integration matures over approximately six weeks, the repair was biomechanically sound from the beginning. This allows us to conclude that the prolonged, multi-month recovery period, characterized by a complex two-phase neural reorganization, was not limited by the tendon’s force-bearing capacity but rather reflects the significant challenge of a purely neural learning process.

### The CNS Defaults to a Modular Strategy, Leading to a Maladaptive Conflict

Our primary finding provides a direct answer to the question posed in the introduction: the CNS adapted to the tendon transfer not by developing fine-grained, fractionated control, but by implementing a modular strategy by repurposing entire co-activation modules (Overduin et al., 2008; Berger et al., 2013; Bizzi and Cheung, 2013; Takei et al., 2017). The initial and most immediate neural change was a wholesale swap of the temporal activation patterns of the primary flexor and extensor synergies (Synergy A and B). The fact that the spatial structure of all four synergies remained remarkably stable throughout the months-long experiment (Fig. 7I, S6), in the face of drastically altered biomechanics, strongly supports the hypothesis that these synergies represent stable, neurally constrained building blocks (d’Avella et al., 2003; Bizzi and Cheung, 2013; Takei et al., 2017). While our baseline analysis (Fig. 7I) confirmed that natural variability is negligible (>0.99 similarity), the specific, transient structural deviations observed post-surgery (e.g., in the compensatory wrist flexor Synergy C) suggest that the CNS retains a limited capacity to fine-tune the internal structure of these modules when driven by strong functional demands. Consequently, the distinct adaptive patterns observed in individual non-transferred muscles (e.g., FCR, PL in Monkey A) likely reflect this task-specific tuning of secondary synergies, rather than a breakdown of modular control.

Our result indicated that this modular approach is likely the default strategy. One potential reason is that modulating pre-existing modules may be computationally simpler and metabolically less costly than developing entirely new, fractionated control patterns (Flash and Sejnowski, 2001; Bizzi and Cheung, 2013). The latter would require extensive synaptic plasticity, potentially involving cortical remapping to selectively uncouple previously co-activated muscles (Kitago and Krakauer, 2013). Alternatively, or perhaps complementarily, the preservation of synergies may reflect inherent constraints on neural plasticity, suggesting that the underlying neural circuits encoding these modules are relatively immutable, even when faced with significant changes in peripheral mechanics (Makin and Krakauer, 2023). Regardless of the underlying reason, be it computational efficiency, constraints on plasticity, or both, the CNS appears to prioritize modulating the activation of existing modules when dealing with acute alterations to the musculoskeletal system.

However, this adherence to established synergy structures created a fundamental conflict. Activating the original extensor synergy (Synergy B) after the transfer now inevitably co-activated the surgically transferred EDC, which functioned mechanically as a flexor, alongside non-transferred muscles like ED23 and ECU, which remained anatomical extensors. This internal mechanical antagonism appears to be the root cause of the early maladaptive phase. This interpretation is supported by two key lines of evidence from our results. First, the period of severe behavioral impairment, characterized by off-target reaching and inefficient grasping (Fig. 5B, E), occurred precisely when this flawed “swap” strategy was active. Second, the aggregated EMG activity revealed a sustained and significant increase in the total activation of the conflicted Synergy B in Monkey A (Fig. 11B, F), which we interpret as an energetically costly effort to overcome the internal mechanical antagonism. This scenario can be viewed through the lens of optimal control and cost-benefit analysis, where a cost function is minimized (Wolpert, 1997). The initial “swap” strategy, while computationally cheap to select, incurred an unacceptably high operational cost in terms of both poor task performance (high error) and what was likely excessive energy expenditure. We hypothesize that this profoundly unfavorable cost-benefit ratio likely served as the critical error signal that drove the CNS to abandon this initial strategy.

### Resolution Through Slower Compensatory Adaptations

The CNS did not persist in the failed swap-based strategy. The high metabolic and computational cost of activating a mechanically conflicted synergy likely triggered the second rapid adaptation: the “switch-back” of synergy activation timings toward their original patterns (Fig. 7, 9). This rapid reversion, occurring over just a few days, is characteristic of an error-based learning mechanism, a form of adaptation that is profoundly impaired by cerebellar damage across a range of tasks, including adaptation to force fields (Smith and Shadmehr, 2005), prismatic shifts (Martin et al., 1996), and split-belt treadmills (Morton and Bastian, 2006). It is plausible that the CNS operates with an implicit threshold for an acceptable cost/performance ratio; once the persistent task failure and high muscular co-contraction of the swap strategy exceeded this threshold, a swift recalibration was initiated. This reversion, despite the remaining mechanical antagonism from the transferred tendons, represents a “good enough” solution where functional success is prioritized over perfect efficiency (Mussa-Ivaldi and Bizzi, 1995; Ranganathan et al., 2013; Gijsberts et al., 2014). Our quantitative analysis supports this interpretation: while the CNS successfully restored the temporal structure of the motor commands (high cosine similarity), it did not perfectly replicate the pre-surgery state. As shown in Figure S8, significant differences in activation amplitude persisted in specific phases of the movement, and permutation tests confirmed the profiles remained statistically distinct. This suggests the system settled into a stable, functional attractor that was sufficiently close to the original manifold to execute the task, without expending the computational or metabolic cost required for a perfect restoration. The rapid timescale of this change is highly consistent with the cerebellum’s proposed role as a forward model, predicting the sensory consequences of motor commands and driving rapid learning in response to sensory prediction errors (Popa et al., 2016). Our lag analysis (Fig. S2) confirmed that the timing of motor commands fluctuated significantly before stabilizing.

Crucially, this switch-back to a less conflicted state, presumably representing the involvement of the error-based learning process, was likely enabled only by the concurrent development of slower, compensatory strategies that provided an alternative means to achieve the task goal. The primary compensation was a learned use of the tenodesis effect. Over weeks, the monkeys gradually increased the activation of the wrist flexor synergy (Synergy C) during hand opening. This wrist flexion biomechanically generates passive finger extension (Zajac, 1992; Cash and Jones, 2011), providing a viable new method for hand pre-shaping. This learned, compensatory behavior, confirmed by kinematic analysis (Fig. 12), ultimately allowed the CNS to abandon the maladaptive synergy swap. This gradual, exploratory process is distinct from the following rapid adaptation and represents a form of motor skill acquisition, a process which may be associated with plasticity in cortical structures like the motor cortex and basal ganglia. (Kitago and Krakauer, 2013).

### A Multi-Timescale Model Reconciles Smooth Behavioral Recovery and Abrupt Neural Reorganization

The apparent paradox between the smooth, gradual recovery of motor function and the abrupt, switch-like reorganization of the underlying neural control strategy suggests that adaptation is not a single process. Instead, we propose it results from the interaction of at least two distinct adaptive systems operating in parallel on different timescales, a concept aligned with established two-state models of motor learning (Smith et al., 2006a; Trewartha et al., 2014).

We map the gradual acquisition of the compensatory tenodesis strategy to the slow process of this model. This iterative component is responsible for building the robust, stable motor skills necessary for long-term recovery. In contrast, we propose that the fast process drives the initial ‘swap’ strategy and its subsequent abandonment. This system appears to be sensitive to immediate performance errors and metabolic costs, triggering the rapid ‘switch-back’ to the original synergy timings (recalibration) once the initial strategy proves inefficient.

The critical insight from our data lies in the interaction between these timescales. We propose that the slow system effectively ‘gates’ the fast system: the abrupt neural reorganization (the switch-back) is likely not an independent event, but a transition enabled only when the slow learning (tenodesis) reaches a “good enough” threshold to support function. This hierarchical interaction resolves the tension between the smooth behavioral curve and the sharp neural transition, ensuring a stable long-term motor plan (Fig. 14).

**Figure 14:**
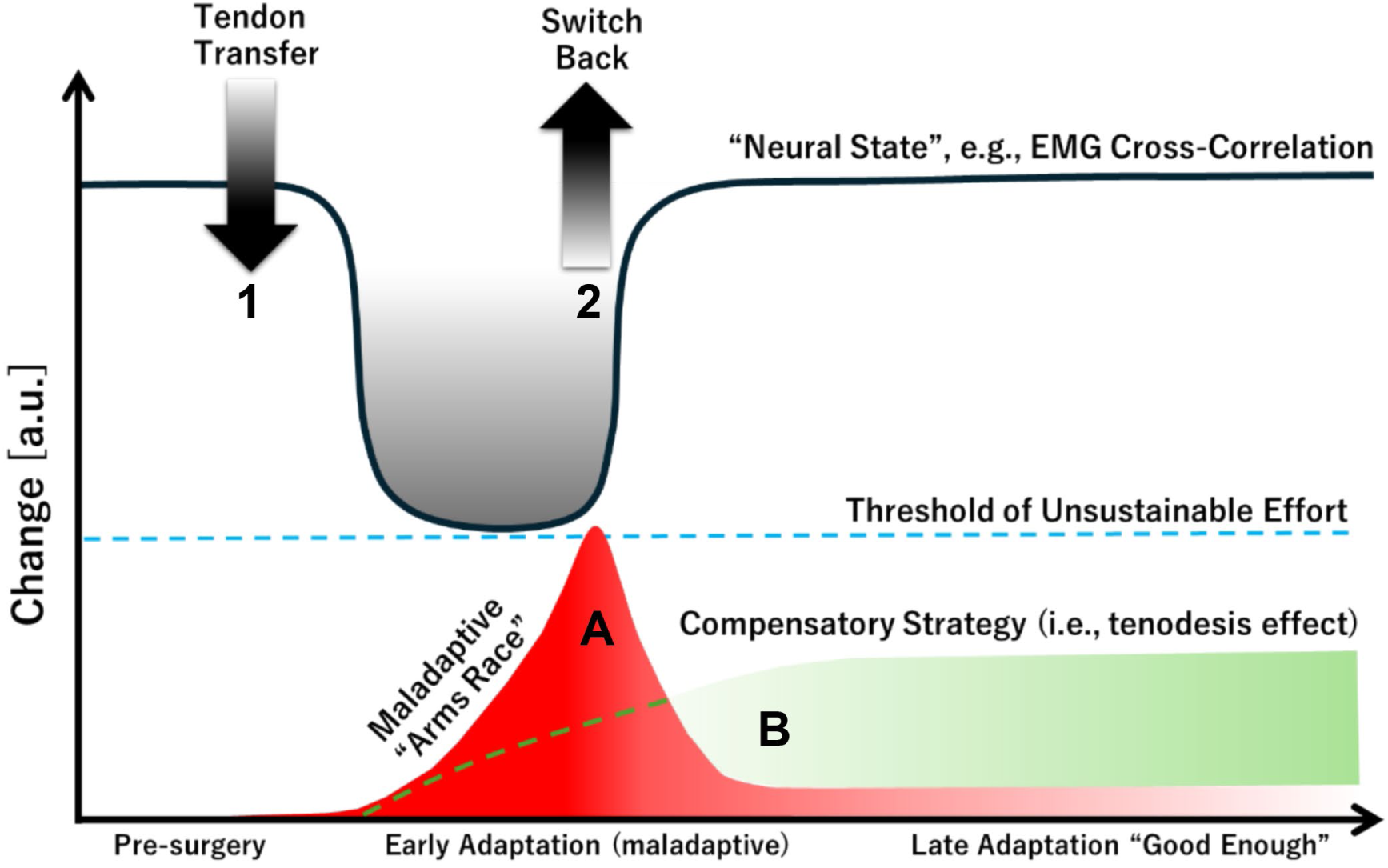
A Proposed Model of Multi-Timescale Adaptation Following Tendon Transfer. This schematic illustrates the hypothesized interaction between fast and slow adaptive processes driving recovery. The initial Tendon Transfer triggers a rapid but maladaptive ‘swap’ of motor commands (Fast Adaptation 1), leading to a maladaptive state. During this phase, two slower processes are hypothesized to occur in parallel: a costly ‘arms race’ within the conflicted synergy (Slow Process A, red curve) and the gradual development of a functional compensatory strategy (Slow Process B, green curve). When the ‘arms race’ reaches a threshold of unsustainable cost (dashed blue line), a second Fast Adaptation (’Switch-Back’, 2) is triggered. This allows for the abandonment of the flawed strategy and the adoption of a stable, ‘good enough’ solution, which is now supported by the newly learned compensatory strategy. The gray line represents the observed neural data (e.g., cross-correlation coefficients of electromyography and temporal activation profiles of muscle synergies), which reflects this two-phase process.

### Broader Implications and Limitations

#### Mechanisms of Neural Adaptation

Our findings of a multi-timescale adaptation process involving stable muscle synergies resonate with, yet also extend, previous work on motor learning. Early studies in primates demonstrated a capacity for eventual positive functional readaptation after nerve crossing, moving beyond the initial maladaptive reversals also seen in our monkeys (Sperry, 1947). However, EMG studies in human patients suggest this adaptation may be incomplete, as original ‘old’ muscle activation patterns often persist alongside newly learned ones (Illert et al., 1986). This aligns with our observation of synergy timing modulation rather than structural reorganization, a finding strongly supported by ‘virtual surgery’ studies indicating the CNS preferentially adapts by recombining existing motor modules (Berger et al., 2013).

#### Biomechanical Constraints and Sensory Feedback

The maladaptive conflict (’arms race’) observed during the early phase may be exacerbated by peripheral biomechanical factors. Rodent models show that epimuscular myofascial force transmission can cause transferred muscles to generate unexpected antagonistic forces depending on joint angle (Maas and Huijing, 2012). Such biomechanical conflicts likely contributed to the cost and ultimate abandonment of the initial swap strategy. Furthermore, the initial, rapid synergy swap may be linked to a ‘sensory surprise’ from altered proprioceptive feedback, driving a fast but flawed response (Kitazawa et al., 1995; Martin et al., 1996; Smith et al., 2006b; Edwards et al., 2012; Petitet et al., 2018). This multi-level network reorganization presents a more complex challenge than the gain modulation within single or specific CNS pathways seen in studies of spinal reflex conditioning (Thompson and Wolpaw, 2014). A comprehensive model must ultimately connect the global synergy reorganization with these local, interacting feedback loops, considering factors such as the roles of spinal reflex pathways (Smith et al., 2006c) and fusimotor drive (Vallbo et al., 1979; Hospod et al., 2007).

*Task-Dependent Differences.* The observation that both monkeys, engaged in both a highly controlled learned task and a more naturalistic grasping task, showed similar adaptive patterns strengthens the generalizability of our findings (Berniker et al., 2013). However, the difference in adaptation rates likely reflects the distinct constraints of these tasks. Monkey A’s precision task required exact force vector production, potentially prolonging the neuromuscular conflict. In contrast, Monkey B’s naturalistic task allowed for more flexible kinematic solutions, such as the tenodesis grasp, to be discovered and employed more rapidly. This suggests that while the core neural mechanism (swap-and-revert) is conserved, the timescale of its resolution is sensitive to task constraints.

#### Motor Memory and Unlearning

Our paradigm raises critical questions about the distinction between unlearning a maladaptive strategy and relearning a functional one (Kitago et al., 2013). The disappearance of the initial “swap” strategy could be due to passive decay, active inhibition (extinction), or interference from a new, stronger memory (Villalta et al., 2013). The literature increasingly suggests that unlearning is an active process, and that the new, successful compensatory strategy likely supplants the maladaptive one through interference. Our results lend weight to this view, suggesting functional recovery is a process of competitive memory formation.

#### Future Directions

A valuable future direction would be to combine the empirical findings of this study with theoretical biomechanical modeling. A subject-specific musculoskeletal model could provide a more precise mapping of how the tendon transfer altered the biomechanical plant. Integrating these empirical and theoretical approaches represents a critical next step in understanding the complex interplay between biomechanics and neural adaptation (Nakajima et al., 2023).

### Implications for Sensorimotor Rehabilitation

These findings may hold implications for sensorimotor rehabilitation. The observation of a distinct maladaptive phase followed by compensation and recovery suggests a staged approach could be beneficial (Hatem et al., 2016; Kakavas et al., 2025). An early phase of therapy might focus on minimizing sensory prediction errors caused by the altered proprioceptive feedback, and reducing maladaptive responses through techniques aimed at integrating the altered body schema (e.g., using visual feedback or task-oriented practice) (Nudo et al., 1996; Takeuchi and Izumi, 2012; Zeiler and Krakauer, 2013; Foell et al., 2014; Levin et al., 2015; De Nunzio et al., 2018; Sattin et al., 2023; van Vliet et al., 2023). This stage would guide the fast, error-driven learning system. A later phase could then employ task-specific training to promote effective compensatory strategies, facilitating the abandonment of maladaptive patterns and the consolidation of a stable motor strategy (Murata et al., 2008; Maier et al., 2019). This second stage engages the slow, skill-acquisition system by creating a rich, problem-solving environment. Finally, a third stage would focus on intensive, high-repetition practice to automate the new functional motor plan, making it robust for real-world use through dosage and generalization (Winterbottom and Nilsen, 2024). This neurobiologically-informed staging, guiding fast adaptation, then facilitating slow skill learning, then driving consolidation, may offer a more logical and effective path to functional recovery.

## METHODS

### Animals

Data were collected using two male macaque monkeys (*Macaca fuscata*; Monkey A: 7.8 kg and Monkey B: 9.9 kg) purpose-bred at the National Bioresource Project (NBRP). They were kept in custom-made primate cages, allowing for potential pair housing. Both monkeys were trained to perform a simple grasping task. During experimental recordings, the monkeys were seated in a primate chair without further restraint. Their head movement remained unrestricted at all times.

### Procedures

All procedures were designed to minimize discomfort and pain and approved by the institutional Animal Care and Use Committees at the National Center of Neurology and Psychiatry (NCNP), Tokyo, Japan. Details of the surgical operations, experimental setup, and procedures for electromyography (EMG) recordings have been previously described (Takei and Seki, 2010).

### EMG implant surgery

Both animals were familiarized with the experimental setup and trained to perform the behavioral task prior to surgery. After an initial training period, EMG electrodes were chronically implanted subcutaneously into several muscles of the left forearm including the FDS, FDP, and EDC (Figs. 1, 3C, and 4C). Individual muscles were localized and confirmed using electrical microstimulation before implanting EMG electrodes chronically by two methods. For muscles involved in the crossed tendon transfer procedure, muscle fascia was cut and EMG wires with looped ends were placed on top of the muscle belly. These were then secured by reclosing the fascia with absorbable suture threaded through the loop (AS-361, stainless-steel Cooner wire, Conner Wire Co., Chatsworth, CA, USA). All other muscles were directly implanted into the muscle belly using a 22-gauge hypodermic needle. Each wire was threaded into the needle tip, folded back along the shaft, and inserted into the muscle before carefully retracting the needle. Electrode separation for bipolar recordings was approximately 5–10 mm. Note that some EMG recording sites were lost over time. Surgical procedures were carried out under deep general anesthesia (sevoflurane 1.5 to 2.5% in 2:1 O2/N2O) and with full aseptic precautions. Heart rate, blood pressure, body temperature, and blood oxygen saturation were monitored throughout surgery. Analgesics and antibiotics were administered intramuscularly for at least one week postoperatively.

### Tendon-Transfer Surgery

After a minimum four-week recovery period from EMG implant surgery, TT surgery was performed. The FDS and EDC tendons were cut as distally as possible (immediately below and above the wrist for FDS and EDC, respectively) to avoid damaging the Golgi tendon organ located near the junction between the muscle fibers and tendon (Schoultz and Swett, 1972; Jami, 1992). Tendons were then guided either through the gap between the radius and ulnar bone (Monkey A), or around the wrist and then re-attached to the tendon of the antagonist muscle using a tendon graft harvested from the plantaris tendon of the lower limb (Monkey B; EDC → FDS: direct connection; and FDS → EDC: tendon grafting). The tendon coaptation was performed using a Pulvertaft weave technique (weaves > 2) with high-strength non-absorbable suture material to ensure immediate and robust mechanical stability post-surgery (Graham et al., 2023). This was only necessary in Monkey B since the tendons used for the cross-transfer were too short. This surgical procedure aimed to reverse the primary mechanical actions of the manipulated muscles, making the FDS tendon an effective finger extensor and the EDC tendon an effective finger flexor. The success and nature of this mechanical rearrangement were verified post-operatively via direct muscle stimulation (see Methods subsection ‘Tendon transfer confirmation’ and Fig. 2). The general organization and function of the major forearm muscles involved in finger flexion and extension in *Macaca fuscata* are broadly comparable to those in humans (Vanhoof et al., 2021; Yan et al., 2022).

Surgical procedures were carried out under deep general anesthesia (sevoflurane 1.5 to 2.5% in 2:1 O_2_/N_2_O) and with full aseptic precautions. Heart rate, blood pressure, body temperature, and blood oxygen saturation were monitored throughout surgery. Analgesics and antibiotics were administered intramuscularly for at least one week postoperatively. Both monkeys wore a plaster cast post-surgery, but its effectiveness was limited (lasting approximately one week).

### Tendon transfer confirmation

To verify the success and long-term stability of TT surgery, two procedures (Fig. 2) were implemented. During the first procedure, the monkey was sedated and its arm was secured to a metal frame (Fig.2A). Electrical stimulation was then applied to either the FDS or EDC muscle (50 mA, DS8R, Digitimer, Welwyn Garden City, UK), while ultrasound scans of tendon movements were concurrently captured (SONIMAGE MX1, Konica Minolta, Inc., Tokyo, Japan (Nordez et al., 2009; Dieterich et al., 2017).

Figure 2B shows a sonogram of the FDS muscle and its intramuscular tendons. The left side displays a static image of the FDS muscle at a specific moment, with white arrows indicating the FDS tendon used for measurement. The right side shows staggered images (top white box) of the FDS tendon displacement triggered by muscle stimulation. The lower right inset illustrates the measurement area. The displacement wave area represents the intensity of muscle contraction, and was calculated by measuring the average duration (a, in seconds) and amplitude (b, in cm) of three successive waves. The area (a*b/2) and regression line for FDS (red) and EDC (blue) (R^2^ > 0.5 for FDS) for days 0, 7, and 105 post-TT are shown (Fig. 2C). The results suggest that the muscle contractions induced by direct electrical stimulation remained nearly constant.

For the second procedure, high-speed videos (37U Series Color Industrial Camera, The Imaging Source, Charlotte, NC, USA) of the monkey’s finger movements were recorded while electrically stimulating the FDS or EDC. Markers were placed on the nails of the index, middle, and ring fingers (Fig. 2A) to measure finger displacement in xyz-dimensions using DeepLabCut software (Mathis et al., 2018; Nath et al., 2019). The sum of Euclidean distances of each marker from the origin of the three-dimensional (3D) coordinate system was computed as a scalar quantity. Post-surgery, movement along the z-axis reversed, indicating a shift from finger flexion to extension due to TT (Fig. 2D; blue: pre-TT on surgery day, dark brown: post-TT on surgery day, light brown: 1 wk post-TT, and red: 3 wk post-TT). The scalar quantities of each finger did not significantly change during muscle stimulation on days 0-, 7-, and 105-days post-TT (Fig. 2E), suggesting there was no postoperative tendon rupture or slackening.

### Data recordings

EMG signals were recorded using the AlphaLab SnR system (Alpha Omega Engineering, Hamerkava St. 6, Ziporit Industrial Zone, P.O. Box 810, Nof HaGalil (Nazareth Illit) 1789062, Israel), displayed online and then stored on a hard drive for later off-line analysis using MATLAB (MathWorks, Natick, MA, USA). Data were recorded at 11 kHz and processed for analysis as follows. 1. Down-sampled to 5 kHz; 2. 50 Hz high-pass filtered (6th order Butterworth filter); 3. rectified; 4. 20 Hz low-pass filtered (6th order Butterworth filter); and 5. down-sampled to 100 Hz. Behavioral task events were recorded as transistor–transistor logic (TTL) signals using the AlphaLab SnR system. Monkey behavior was further recorded by two cameras (Sanyo VPC-WH1, 60 fps [Sanyo, Osaka, Japan]; and Casio EX-100F, 240 fps [Casio, Tokyo, Japan] for monkeys A and B, respectively) from two different angles (top and side view). The images were later used to detect additional behavioral events (contact time with the object and pull onset for Monkey A; and food touch and food lift onset for Monkey B), and detection of possible changes in the animal’s general movement pattern. Kinovea software was used for this video analysis (free and open-source software; https://www.kinovea.org/).

### Behavioral task

Before and after EMG surgery, the monkeys were trained on a simple grasping task that involved a small object attached to a rod (Fig. 3A for Monkey A; and Fig. 4A for Monkey B). For Monkey A, object 1 was a small rod placed between two side walls encouraging the monkey to grasp the rod with a precision grip using the tips of the index finger and thumb (controlled grasp). The force used to compress the spring while pulling was low and the distance moved short. Object 2 had to be grasped in the same way. However, the force to compress the spring was higher and the distance moved longer (object 1: 300 cN, 4 mm; and object 2: 800 cN, 30 mm). The monkey was asked to grasp and hold the objects for approximately 300–500 ms (Fig. 3A–D). After completing the trial, the monkey was rewarded with a piece of fruit that had to be taken from the experimenter’s palm. The task sequence for Monkey A ranged from object 1 hold onset to object 2 hold offset (Fig. 3C, D; and ‘Data Analysis’ below). For Monkey B, object 1 was a small rod that was grasped and pulled (300 cN, 4 mm) using a lateral prehension grip (like using a door key), and resembling a power grip rather than a natural grasp. However, instead of grasping object 2, Monkey B picked up a piece of fruit from an allocated location. In doing so, the monkey had to cross the path of a photo cell which was detected by the recording system (Fig. 4A–D). The task sequence to be analyzed for Monkey B ranged from object 1 hold onset to food touch, which was indicated by a LED (Fig. 4C, D; and ‘Data Analysis’ below).

For Monkey A, the task sequence was as follows (Fig. 3B; and Supplementary Video S1). From a starting position in front of the monkey, it had to lift its arm and move towards object 1, grasp, pull, and then hold the object for 500 ms. Immediately after releasing object 1, Monkey A proceeded to move its hand towards object 2, which was located to the right of object 1. Again, the monkey had to grasp, pull, and then hold the object for another 500 ms. Each hold period was accompanied by an audio signal which stopped once the hold duration was sufficient. After completing the task, which was signaled by another acoustic signal, the monkey was rewarded with a piece of fruit presented by the experimenter and taken by the monkey.

For Monkey B, the first part of the sequence was identical. However, after releasing object 1, Monkey B was required to pick up a piece of fruit from an allocated location (Fig. 4A, B; and Supplementary Video S5). In doing so, the monkey passed a photocell in front of the food well. This event was detected by the recording system and a video camera (a red LED triggered by activation of the photocell). For the analysis, only object 1 and the food grasp were considered in monkeys A and B, respectively.

### Data analysis

#### EMG analysis

EMG data were normalized to the average time for each monkey to complete a trial (object 1 hold onset ➔ object 2 hold offset/LED offset = 100%). Data were cropped and aligned according to the time stamps (Obj1 hold offset ± 15%; and LED onset [food touch] ± 15% for monkeys A and B, respectively). Recorded EMGs from the pre- and post-tendon surgery period were averaged for each recording and compared over experimental sessions. For Monkey A, FDS and EDC were analyzed both pre- and post-TT. For Monkey B, FDS and EDC were analyzed pre-TT, but only EDC was analyzed post-TT due to signal deterioration.

Both monkeys required time to recover from surgery. Post-TT recordings were resumed once the monkeys were able to perform the task independently (i.e., without assistance from the experimenter) and met specific trial count criteria (starting 29 and 20 days post-surgery for monkeys A and B, respectively). Post-operative recording sessions were conducted frequently throughout the recovery period, though not at fixed daily intervals, reflecting the practical constraints of long-term behavioral experiments (e.g., animal cooperation, experimental scheduling) and the aim to capture data during key phases of adaptation. Within any given session included in the analysis, behavioral (video) and EMG data were collected concurrently. To be included in the behavioral analyses (e.g., grip formation times, off-target reaching, kinematics), a session required a minimum of 20 successful trials. For the more demanding muscle synergy analyses, a higher threshold of at least 100 successful trials was required to ensure robust factorization. This difference in criteria may have resulted in slightly different sets of recording days being represented across behavioral versus synergy-related figures. After tendon surgery, EMG signals for FDS, FCU, and FCR deteriorated in Monkey B. Therefore, there is no experimental data for these muscles in this monkey and they were excluded from the muscle synergy analysis.

#### Synergy analysis

The EMG envelopes obtained from pre-processing raw EMG data were divided by the mean value to normalize activity. Muscle synergies were then extracted for each session using NMF (d’Avella et al., 2003). NMF decomposes the EMG data matrix M, as a product of two matrices C and W:

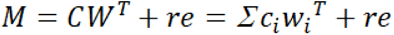

where the vector wi and column of matrix W are muscle synergies; the vector ci, and column of matrix C are their temporal coefficients; and re represents the residuals. Multiplicative update rules were used for decomposition (Lee and Seung, 1999). Updating matrices for decomposition from 20 different random initial matrices was started, and factorization results with least residuals among the 20 results were used. To improve robustness of the muscle synergies, one set of muscle synergies was extracted from EMG data of multiple trials of each day. A k-fold cross validation was used, with k = 4 i.e., EMG data set of multiple trials during the day was split into four data sets, in which three data sets were for training and one data set was for testing. Synergy matrix W was calculated from three training data sets using NMF, and then a coefficient matrix C was calculated by decomposing the EMG matrix M of the test data set with a fixed W matrix of the training data set. This operation was repeated by changing the test data set for each four data sets. Finally, the extracted four C matrices and four W matrices were averaged to obtain daily synergies.

The number of synergies was determined as the number where VAF exceeded a threshold. This threshold was set as 0.8. To clarify the effect of TT, the number of synergies for each monkey was determined, assuming that trial-by-trial variation in synergies within the same monkey was small. MATLAB ‘nnmf’ function was used for NMF.

To determine whether the weights of muscle synergies changed after TT, the following procedure was used. First, the average of each pre-TT spatial synergy was calculated and used as control data to calculate the cosine distance for all post-TT spatial synergies. This generated four distance relationships for each pre-TT synergy. If the change in spatial synergy before and after TT was small, then only one of the four cosine distances (e.g., synergy A before TT and synergy A after TT) should be significantly smaller. Two-way ANOVA was conducted using the type of synergy and session as factors for the cosine distance. The significance of synergy pairs was identified using the Bonferroni post-hoc test.

#### Quantitative Comparison of Synergy Profiles

To quantify the degree of ‘reversion’ in the late adaptation phase, we compared the synergy activation profiles from the pre-surgery period with those from the final recording day using three distinct metrics. First, Cosine Similarity was calculated to assess the similarity in the shape of the temporal profiles independent of amplitude. Second, a Permutation Test (n=10,000 iterations) was performed to test whether the specific trajectory of the post-surgery profile was statistically distinguishable from the pre-surgery distribution. Third, to identify specific phases of the task where activation differences persisted, we conducted a point-by-point Wilcoxon rank-sum test at each time point of the normalized task cycle, applying a Bonferroni correction for multiple comparisons.

#### Cross-correlation analysis

Cross-correlation analysis was performed to examine the similarity between pre-and post-surgery EMG signals as well as temporal activation profiles of extracted muscle synergies. For the EMG signal analysis presented in Figure 6, the cross-correlation coefficient at zero-time lag was calculated. For the synergy analysis presented in the main figures (Figs. 9, 10), the cross-correlation coefficient at zero-time lag was calculated using MATLAB (MathWorks, corrcoef function) to quantify similarity without temporal shifts. These zero-lag coefficients were plotted over experimental sessions. In a supplementary analysis, we also computed the cross-correlation allowing for variable time lags using MATLAB’s xcorr function (normalized using the ‘coeff’ option). From this, we extracted the specific time lag that yielded the maximum cross-correlation coefficient (optimal lag) for each session for both the muscle synergies (Supplementary Figure S9) and the individual EMG signals (Supplementary Figure S1). In this analysis, a positive lag indicates that the post-surgery activation profile is delayed (occurs later) relative to the pre-surgery baseline, while a negative lag indicates it is advanced (occurs earlier).

Based on cross-correlation analysis, five ‘landmark days’ were identified for further analysis (Fig. 6P). These days represent distinct stages in the recovery process and excluded pre-TT control data. The first landmark day was chosen from one of the initial recording sessions after TT surgery (days 29 and 22 in monkeys A and B, respectively) when the cross-correlation coefficients had changed significantly compared with control data. The second day was then chosen from a time period just before the switch-back, when the cross-correlation coefficients had started to return to their original values (days 64 and 36). Another day was then picked from the period when the coefficients were still changing significantly (days 69 and 44) and before starting to saturate. At this point the next day was defined (days 79 and 48). Finally, the final landmark day was chosen from one of the last recording sessions when the behavior had fully recovered (days 99 and day 64 for monkeys A and B, respectively).

#### Behavioral analysis

To examine behavioral recovery, the duration, onset, and offset of object and food grasps were analyzed. Event times extracted from the video analysis were used for alignment of EMG data and subsequent cross-correlation and synergy analyses, along with recorded event TTL signals. For each experimental session, the first 20 trials of each video recording were analyzed. In Monkey A, the following events were detected and in-between times stored in ms: touch onset, pull onset, and hold onset (resulting in grasp duration and pull time) for object 1 and object 2. In Monkey B, the following events were detected and in-between times stored: touch onset, pull onset, and hold onset (resulting in grasp duration and pull time) for object 1; and for the food grasp component of the task, LED onset, food-touch onset, food-lift onset, LED offset, and movement end (contact of food with mouth).

Contact times (touch onset ➔ pull onset) for Monkey A were plotted in ms (mean ± SD) over number of days from tendon surgery for object 1. For Monkey B, contact times with food (food-touch ➔ food-lift onset) were plotted in ms (mean ± SD).

ANOVA was performed to compare control data recorded before and after tendon surgery. In total, 5 and 3 control sessions vs. 35 and 22 experimental sessions were used for analyses in monkeys A and B, respectively.

To quantify and compare the baseline behavioral variability of these metrics, we analyzed all pre-TT trials for each animal’s respective tasks. We calculated the Coefficient of Variation (CV), defined as the standard deviation divided by the mean, for each metric (grip formation time, pull time, and grasp aperture). To formally test the difference in variance for the grip formation task (Fig. 5A, D), we used the non-parametric Ansari-Bradley test. To quantify the observed off-target reaching behavior, video footage was analyzed and the means of 10 consecutive trials calculated for each session as follows. For Monkey A, time spent within a specific spatial window covering the area behind and between the two objects was measured; the monkey passed through this space almost exclusively while exhibiting the impairment (see Fig. 3E, crossing of the yellow dotted line). In Monkey B, the time spent in contact with the rear plate of the object was measured (see Fig. 4E).

In addition to the primary behavioral metrics, we performed a more detailed video analysis of movement kinematics and performance. For Monkey A, we quantified the ‘pull time’ for each trial, defined as the duration from the moment the monkey started pulling the object to the moment hold onset. For Monkey B, we quantified the ‘grasp aperture’. This was defined as the distance between the tips of the index finger and thumb, measured just before the monkey made contact with the food pellet.

#### Joint kinematics analysis

To determine whether compensatory movements occurred, steady state changes in joint angles were examined before and after recovery. Using DeepLabCut (Mathis et al., 2018; Nath et al., 2019), key points on the fingers and wrists were tracked from experimental videos. From the obtained key points, the vector *v_2_* connecting the wrist and fingers and vector *v_2_* connecting the arm and wrist were estimated (Fig. S10). The joint angle θ was calculated as follows:

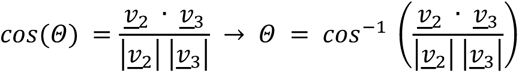

To quantify the learning and refinement of the compensatory tenodesis strategy over time, we analyzed the trial-by-trial relationship between wrist and MCP joint angles for Monkey B. For each recorded day, we performed a linear regression between the wrist angle (predictor) and the MCP angle (outcome) across all successful trials (mean of 20 trials ±SD for each landmark day, taken 83ms before food touch). The strength of the coupling was quantified using the coefficient of determination (R²). These daily scatter plots and their corresponding R² values were visualized to track the evolution of the kinematic coupling from the pre-TT baseline through the post-surgery adaptation period.

## Supporting information

supplementary informations

## DATA, MATERIALS, AND SOFTWARE AVAILABILITY

Matlab scripts and raw data will be made available online. All study data are included in the article and/or SI Appendix.

## ACKNOWLEDGEMENTS

We would like to express our sincere gratitude to Drs. Francisco Valero-Cuevas and Andrea d’Avella for their invaluable comments and constructive suggestions on the earlier version of the manuscript, and Drs Nicholas Schweighofer and Gerald E. Loeb for stimulating discussion. We would also like to thank Dr. Naomichi Ogihara and Prof. Toshiyasu Nakamura for their valuable insights and technical advice on tendon surgery in the hand and forearm of primates. We are grateful to Drs. Kumiko Oida and Chika Sasaki for their medical care of the experimental animals, vital help during surgery, and training of the animals. Lastly, we would like to thank Masahi Koizumi for designing parts of the experimental setup and Drs. Joachim Confais, Shinji Kubota, Saeka Tomatsu, Tatsuya Umeda, and Natsumi Uchida for their support and assistance during the proceeding pilot study. We thank Rachel James, PhD, from Edanz (https://jp.edanz.com/ac) for editing a draft of this manuscript.

This work was supported by Grants-in-Aid for Scientific Research from the Ministry of Education, Culture, Sports, Science, and Technology of Japan (26120003, 26250013, 15K21754, 19H05724, 19H01092, 23H05488, and 24K21313), the Japan Agency for Medical Research and Development (JP24gm0010009), the Japan Science Technology Agency Precursory Research for Embryonic Science and Technology program, commissioned research (no. 22102) from the National Institute of Information and Communications Technology, and commissioned research by National Institute of Information and Communications Technology (NICT), Japan (all to K.S.). This work is also supported in part by the NSF CRCNS Japan-US 2113096 to K.S. (Subaward PI) and Francisco Valero-Cuevas (PI). The content is solely the responsibility of the authors and does not necessarily represent the official views of the NSF.

## REFERENCES

Berger DJ, Gentner R, Edmunds T, Pai DK, d’Avella A (2013) Differences in Adaptation Rates after Virtual Surgeries Provide Direct Evidence for Modularity. The Journal of Neuroscience 33:12384–12394.

Berniker M, O’Brien MK, Kording KP, Ahmed AA (2013) An Examination of the Generalizability of Motor Costs. PLOS ONE 8:e53759.

Bizzi E, Cheung VC (2013) The neural origin of muscle synergies. Frontiers in Computational Neuroscience Volume 7–2013.

Bowlus TH, Lane RD, Stojic AS, Johnston M, Pluto CP, Chan M, Chiaia NL, Rhoades RW (2003) Comparison of reorganization of the somatosensory system in rats that sustained forelimb removal as neonates and as adults. J Comp Neurol 465:335–348.

Cash DJW, Jones JWM (2011) The role of tenodesis in surgery of the upper limb. The Journal of Bone & Joint Surgery British Volume 93-B:285–292.

Cheung VC, Turolla A, Agostini M, Silvoni S, Bennis C, Kasi P, Paganoni S, Bonato P, Bizzi E (2012) Muscle synergy patterns as physiological markers of motor cortical damage. Proc Natl Acad Sci U S A 109:14652–14656.

d’Avella A, Saltiel P, Bizzi E (2003) Combinations of muscle synergies in the construction of a natural motor behavior. Nature Neuroscience 6:300–308.

Davidson PR, Wolpert DM (2003) Motor learning and prediction in a variable environment. Curr Opin Neurobiol 13:232–237.

De Nunzio AM, Schweisfurth MA, Ge N, Falla D, Hahne J, Gödecke K, Petzke F, Siebertz M, Dechent P, Weiss T, Flor H, Graimann B, Aszmann OC, Farina D (2018) Relieving phantom limb pain with multimodal sensory-motor training. J Neural Eng 15:066022.

Dieterich AV, Botter A, Vieira TM, Peolsson A, Petzke F, Davey P, Falla D (2017) Spatial variation and inconsistency between estimates of onset of muscle activation from EMG and ultrasound. Sci Rep 7:42011.

Edwards MJ, Adams RA, Brown H, Pareés I, Friston KJ (2012) A Bayesian account of ‘hysteria’. Brain 135:3495–3512.

Flash T, Sejnowski TJ (2001) Computational approaches to motor control. Curr Opin Neurobiol 11:655–662.

Fleury L, Prablanc C, Priot AE (2019) Do prism and other adaptation paradigms really measure the same processes? Cortex 119:480–496.

Foell J, Bekrater-Bodmann R, Diers M, Flor H (2014) Mirror therapy for phantom limb pain: brain changes and the role of body representation. Eur J Pain 18:729–739.

Gaetz W, Dockstader C, Furlong PL, Amaral S, Vossough A, Schwartz ES, Roberts TPL, Scott Levin L (2023) Somatosensory and motor representations following bilateral transplants of the hands: A 6-year longitudinal case report on the first pediatric bilateral hand transplant patient. Brain Res 1804:148262.

Gardenier J, Garg R, Mudgal C (2020) Upper Extremity Tendon Transfers: A Brief Review of History, Common Applications, and Technical Tips. Indian J Plast Surg 53:177–190.

Gijsberts A, Bohra R, Sierra González D, Werner A, Nowak M, Caputo B, Roa MA, Castellini C (2014) Stable myoelectric control of a hand prosthesis using non-linear incremental learning. Front Neurorobot 8:8.

Graham EM, Oliver JD, Hendrycks R, Maglic D, Mendenhall SD (2023) Alternative Tendon Coaptations to the Pulvertaft Weave Technique: A Systematic Review and Meta-Analysis of Biomechanical Studies. HAND 18:446–455.

Green HJ (1997) Mechanisms of muscle fatigue in intense exercise. J Sports Sci 15:247–256.

Hatem SM, Saussez G, Della Faille M, Prist V, Zhang X, Dispa D, Bleyenheuft Y (2016) Rehabilitation of Motor Function after Stroke: A Multiple Systematic Review Focused on Techniques to Stimulate Upper Extremity Recovery. Front Hum Neurosci 10:442.

Hoffmann G, Kamper DG, Kahn JH, Rymer WZ, Schmit BD (2009) Modulation of stretch reflexes of the finger flexors by sensory feedback from the proximal upper limb poststroke. J Neurophysiol 102:1420–1429.

Hospod V, Aimonetti J-M, Roll J-P, Ribot-Ciscar E (2007) Changes in Human Muscle Spindle Sensitivity during a Proprioceptive Attention Task. The Journal of Neuroscience 27:5172–5178.

Hunter DJ, Eckstein F (2009) Exercise and osteoarthritis. J Anat 214:197–207.

Illert M, Trauner M, Weller E, Wiedemann E (1986) Forearm muscles of man can reverse their function after tendon transfers: An electromyographic study. Neuroscience Letters 67:129–134.

Jami L (1992) Golgi tendon organs in mammalian skeletal muscle: functional properties and central actions. Physiol Rev 72:623–666.

Kakavas G, Brilakis E, Papatzikou M, Malliaropoulos N, Mazeas J, Forelli F (2025) Reverse Linear Neuro Periodization Model for Rehabilitation After Arthroscopic Rotator Cuff Repair: A Narrative Review. Clinics and Practice 15:105.

Kitago T, Krakauer JW (2013) Motor learning principles for neurorehabilitation. Handb Clin Neurol 110:93–103.

Kitago T, Ryan SL, Mazzoni P, Krakauer JW, Haith AM (2013) Unlearning versus savings in visuomotor adaptation: comparing effects of washout, passage of time, and removal of errors on motor memory. Front Hum Neurosci 7:307.

Kitazawa S, Kohno T, Uka T (1995) Effects of delayed visual information on the rate and amount of prism adaptation in the human. J Neurosci 15:7644–7652.

Lee DD, Seung HS (1999) Learning the parts of objects by non-negative matrix factorization. Nature 401:788–791.

Levin MF, Weiss PL, Keshner EA (2015) Emergence of virtual reality as a tool for upper limb rehabilitation: incorporation of motor control and motor learning principles. Phys Ther 95:415–425.

Loeb GE (1999) Asymmetry of hindlimb muscle activity and cutaneous reflexes after tendon transfers in kittens. J Neurophysiol 82:3392–3405.

Luauté J, Schwartz S, Rossetti Y, Spiridon M, Rode G, Boisson D, Vuilleumier P (2009) Dynamic changes in brain activity during prism adaptation. J Neurosci 29:169–178.

Maas H, Huijing PA (2012) Mechanical effect of rat flexor carpi ulnaris muscle after tendon transfer: does it generate a wrist extension moment? Journal of Applied Physiology 112:607–614.

Maier M, Ballester BR, Verschure PFMJ (2019) Principles of Neurorehabilitation After Stroke Based on Motor Learning and Brain Plasticity Mechanisms. Frontiers in Systems Neuroscience Volume 13–2019.

Makin TR, Krakauer JW (2023) Against cortical reorganisation. eLife 12:e84716.

Martin TA, Keating JG, Goodkin HP, Bastian AJ, Thach WT (1996) Throwing while looking through prisms. II. Specificity and storage of multiple gaze-throw calibrations. Brain 119 ( Pt 4):1199–1211.

Mathis A, Mamidanna P, Cury KM, Abe T, Murthy VN, Mathis MW, Bethge M (2018) DeepLabCut: markerless pose estimation of user-defined body parts with deep learning. Nature Neuroscience 21:1281–1289.

Mercuri E, Muntoni F (2013) Muscular dystrophies. Lancet 381:845–860.

Morton SM, Bastian AJ (2006) Cerebellar contributions to locomotor adaptations during splitbelt treadmill walking. J Neurosci 26:9107–9116.

Murata Y, Higo N, Oishi T, Yamashita A, Matsuda K, Hayashi M, Yamane S (2008) Effects of motor training on the recovery of manual dexterity after primary motor cortex lesion in macaque monkeys. J Neurophysiol 99:773–786.

Mussa-Ivaldi FA, Bizzi E (1995) The Modular Organization of Motor Control: What Frogs Can Teach Us About Adaptive Learning. IFAC Proceedings Volumes 28:413–418.

Nakajima N, Wang S, Ogihara N, Oya T, Seki K, Funato T (2023) Upper Limb Musculoskeletal Model of Macaque Monkey for Approaching Adaptation Mechanism to Tendon Transfer. Neuroscience 2023 Abstracts, Washington, DC: Society for Neuroscience.

Nath T, Mathis A, Chen AC, Patel A, Bethge M, Mathis MW (2019) Using DeepLabCut for 3D markerless pose estimation across species and behaviors. Nature Protocols 14:2152–2176.

Nordez A, Gallot T, Catheline S, Guével A, Cornu C, Hug F (2009) Electromechanical delay revisited using very high frame rate ultrasound. J Appl Physiol (1985) 106:1970–1975.

Nudo RJ, Wise BM, SiFuentes F, Milliken GW (1996) Neural substrates for the effects of rehabilitative training on motor recovery after ischemic infarct. Science 272:1791–1794.

Osu R, Franklin DW, Kato H, Gomi H, Domen K, Yoshioka T, Kawato M (2002) Short- and long-term changes in joint co-contraction associated with motor learning as revealed from surface EMG. J Neurophysiol 88:991–1004.

Overduin SA, d’Avella A, Roh J, Bizzi E (2008) Modulation of Muscle Synergy Recruitment in Primate Grasping. The Journal of Neuroscience 28:880-892.

Petitet P, O’Reilly JX, O’Shea J (2018) Towards a neuro-computational account of prism adaptation. Neuropsychologia 115:188–203.

Popa LS, Streng ML, Hewitt AL, Ebner TJ (2016) The Errors of Our Ways: Understanding Error Representations in Cerebellar-Dependent Motor Learning. Cerebellum 15:93–103.

Power JD, Schlaggar BL (2017) Neural plasticity across the lifespan. Wiley Interdiscip Rev Dev Biol 6.

Ranganathan R, Adewuyi A, Mussa-Ivaldi FA (2013) Learning to be lazy: exploiting redundancy in a novel task to minimize movement-related effort. J Neurosci 33:2754–2760.

Sattin D, Parma C, Lunetta C, Zulueta A, Lanzone J, Giani L, Vassallo M, Picozzi M, Parati EA (2023) An Overview of the Body Schema and Body Image: Theoretical Models, Methodological Settings and Pitfalls for Rehabilitation of Persons with Neurological Disorders. Brain Sci 13.

Schärli A, Hecht H, Mast FW, Hossner EJ (2024) How spotting technique affects dizziness and postural stability after full-body rotations in dancers. Hum Mov Sci 95:103211.

Schoultz TW, Swett JE (1972) The fine structure of the Golgi tendon organ. J Neurocytol 1:1–26.

Slawinska U, Kasicki S (2002) Altered Electromyographic Activity Pattern of Rat Soleus Muscle Transposed into the Bed of Antagonist Muscle. The Journal of Neuroscience 22:5808-5812.

Smith MA, Shadmehr R (2005) Intact ability to learn internal models of arm dynamics in Huntington’s disease but not cerebellar degeneration. J Neurophysiol 93:2809–2821.

Smith MA, Ghazizadeh A, Shadmehr R (2006a) Interacting Adaptive Processes with Different Timescales Underlie Short-Term Motor Learning. PLOS Biology 4:e179.

Smith MA, Ghazizadeh A, Shadmehr R (2006b) Interacting adaptive processes with different timescales underlie short-term motor learning. PLoS Biol 4:e179.

Smith SA, Mitchell JH, Garry MG (2006c) The mammalian exercise pressor reflex in health and disease. Exp Physiol 91:89–102.

Sperry RW (1940) The functional results of muscle transposition in the hind limb of the rat. Journal of Comparative Neurology 73:379–404.

Sperry RW (1942) Transplantation of motor nerves and muscles in the forelimb of the rat. Journal of Comparative Neurology 76.

Sperry RW (1947) Effect of crossing nerves to antagonistic limb muscles in the monkey. Arch Neurol Psychiatry 58:452–473.

Sugita Y (1996) Global plasticity in adult visual cortex following reversal of visual input. Nature 380:523–526.

Takei T, Seki K (2010) Spinal interneurons facilitate coactivation of hand muscles during a precision grip task in monkeys. J Neurosci 30:17041–17050.

Takei T, Confais J, Tomatsu S, Oya T, Seki K (2017) Neural basis for hand muscle synergies in the primate spinal cord. Proceedings of the National Academy of Sciences 114:8643–8648.

Takeuchi N, Izumi S (2012) Maladaptive plasticity for motor recovery after stroke: mechanisms and approaches. Neural Plast 2012:359728.

Tannenbaum J, Bennett BT (2015) Russell and Burch’s 3Rs then and now: the need for clarity in definition and purpose. J Am Assoc Lab Anim Sci 54:120–132.

Thompson AK, Wolpaw JR (2014) Operant conditioning of spinal reflexes: from basic science to clinical therapy. Front Integr Neurosci 8:25.

Thoroughman KA, Shadmehr R (1999) Electromyographic correlates of learning an internal model of reaching movements. J Neurosci 19:8573–8588.

Trewartha KM, Garcia A, Wolpert DM, Flanagan JR (2014) Fast but fleeting: adaptive motor learning processes associated with aging and cognitive decline. J Neurosci 34:13411–13421.

Valero-Cuevas FJ, Hentz VR (2002) Releasing the A3 pulley and leaving flexor superficialis intact increases pinch force following the Zancolli lasso procedures to prevent claw deformity in the intrinsic palsied finger. J Orthop Res 20:902–909.

Vallbo AB, Hagbarth KE, Torebjörk HE, Wallin BG (1979) Somatosensory, proprioceptive, and sympathetic activity in human peripheral nerves. Physiol Rev 59:919–957.

van Vliet P, Carey LM, Turton A, Kwakkel G, Palazzi K, Oldmeadow C, Searles A, Lavis H, Middleton S, Galloway M, Dimech-Betancourt B, O’Keefe S, Tavener M (2023) Task-specific training versus usual care to improve upper limb function after stroke: the “Task-AT Home” randomised controlled trial protocol. Frontiers in Neurology 14.

Vanhoof MJM, van Leeuwen T, Galletta L, Vereecke EE (2021) The forearm and hand musculature of semi-terrestrial rhesus macaques (Macaca mulatta) and arboreal gibbons (fam.Hylobatidae). Part II. Quantitative analysis. Journal of Anatomy 238:321–337.

Villalta JI, Landi SM, Fló A, Della-Maggiore V (2013) Extinction Interferes with the Retrieval of Visuomotor Memories Through a Mechanism Involving the Sensorimotor Cortex. Cerebral Cortex 25:1535–1543.

Walker JG, Jackson HJ, Littlejohn GO (2004) Models of adjustment to chronic illness: using the example of rheumatoid arthritis. Clin Psychol Rev 24:461–488.

Wester K, Hove LM, Barndon R, Craven AR, Hugdahl K (2018) Cortical Plasticity After Surgical Tendon Transfer in Tetraplegics. Front Hum Neurosci 12:234.

Winterbottom L, Nilsen DM (2024) Motor Learning Following Stroke: Mechanisms of Learning and Techniques to Augment Neuroplasticity. Phys Med Rehabil Clin N Am 35:277–291.

Wolpert DM (1997) Computational approaches to motor control. Trends Cogn Sci 1:209–216.

Yan Y, Sobinov AR, Bensmaia SJ (2022) Prehension kinematics in humans and macaques. J Neurophysiol 127:1669–1678.

Yumiya H, Larsen KD, Asanuma H (1979) Motor readjustment and input-output relationship of motor cortex following cross-connection of forearm muscles in cats. Brain Research 177:566–570.

Zajac FE (1992) How musculotendon architecture and joint geometry affect the capacity of muscles to move and exert force on objects: a review with application to arm and forearm tendon transfer design. J Hand Surg Am 17:799–804.

Zeiler SR, Krakauer JW (2013) The interaction between training and plasticity in the poststroke brain. Curr Opin Neurol 26:609–616.

Zemková E (2022) Physiological Mechanisms of Exercise and Its Effects on Postural Sway: Does Sport Make a Difference? Front Physiol 13:792875.

