## supplementary informations for "Multi-timescale neural adaptation underlying long-term musculoskeletal reorganization"

**(Short title:** Multi-timescale neural adaptation**)**

### Extended data

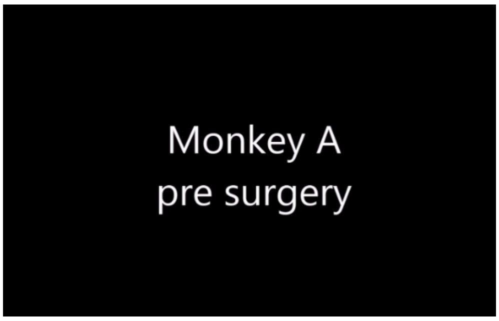


**Supplementary Video S1. Control Behavior (Monkey A – Pre-TT)**

This video demonstrates monkey A's baseline performance on the reaching task before the crossed tendon transfer procedure. It illustrates the typical, coordinated movements exhibited by the monkey in its unaltered state, serving as a control for comparison with post-surgical behavior. Observe the smooth and accurate reaching and grasping motions as the monkey performs the task.

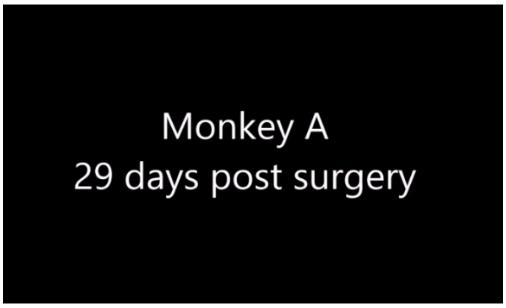


**Supplementary Video S2. 29 Days Post Crossed Tendon Transfer (Monkey A)**

This video documents monkey A's attempts to perform the reaching task 29 days after undergoing the crossed tendon transfer procedure. At this early-stage post-surgery, the video clearly shows the significant impact of the procedure on the monkey's motor control. Observe the marked malcoordination in the monkey's reaching movements. The reaching attempts are furthermore characterized by ‘explorative’ finger movements over the object. The monkey's reliance on the experimenter for support highlights the difficulty it experiences in performing the task independently.

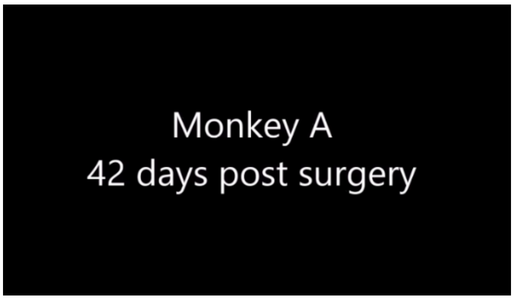


**Supplementary Video S3. 42 Days Post Crossed Tendon Transfer (Monkey A)**

This video shows monkey A's progress 42 days post-surgery. While an improvement in motor control is evident compared to the 29-day mark, the monkey still exhibits some residual deficits. Observe the monkey's attempts to perform the task autonomously. Although it can now perform the task fully without experimenter assistance, it continues to rely on the support of its unaffected arm, suggesting ongoing challenges with coordination and strength.

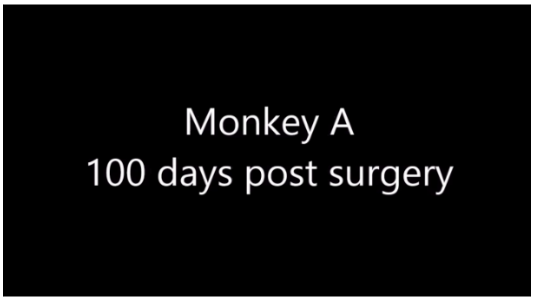


**Supplementary Video S4. 100 Days Post Crossed Tendon Transfer (Monkey A)**

This video demonstrates monkey A's performance 100 days after the crossed tendon transfer. At this point, the monkey has achieved substantial recovery and performs the reaching task with near-normal proficiency. Observe the smooth, coordinated movements, and the monkey's ability to execute the task independently and accurately. This footage showcases the significant recovery of motor function following the procedure.

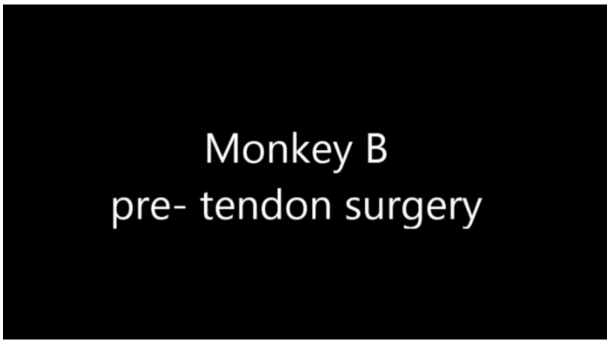


**Supplementary Video S5. Control Behavior (Monkey B - Pre-TT)**

This video establishes the baseline performance for monkey B prior to the crossed tendon transfer procedure. It shows the monkey's typical, coordinated reaching behavior in its unaltered state, providing a control for comparison with its post-surgical performance. Observe the precision and fluidity of the monkey's movements as it executes the task.

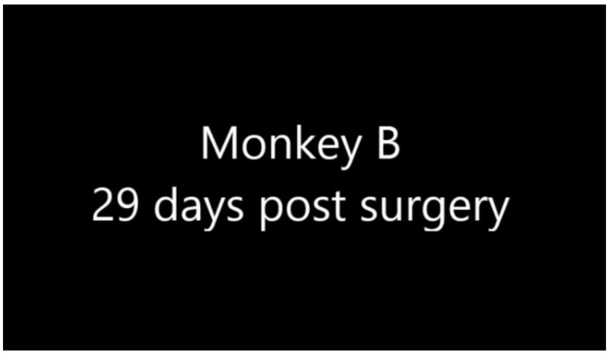


**Supplementary Video S6. 29 Days Post Crossed Tendon Transfer (Monkey B)**

This video documents monkey B's performance 29 days after the crossed tendon transfer. Like monkey A, monkey B exhibits significant motor deficits. The video clearly demonstrates the malcoordinated reaching, characterized by overshooting the target and bumping into objects.

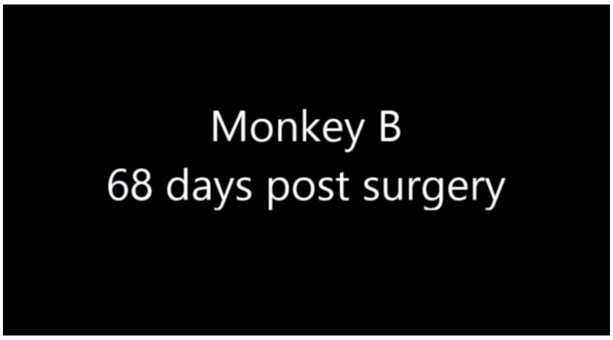


**Supplementary Video S7. 68 Days Post Crossed Tendon Transfer (Monkey B)**

This video shows monkey B's recovery 68 days post-surgery. At this time point, monkey B has achieved a full recovery and performs the reaching task with accuracy and coordination comparable to its pre-surgical baseline. Observe the smooth and efficient movements, demonstrating the successful recovery of motor function.

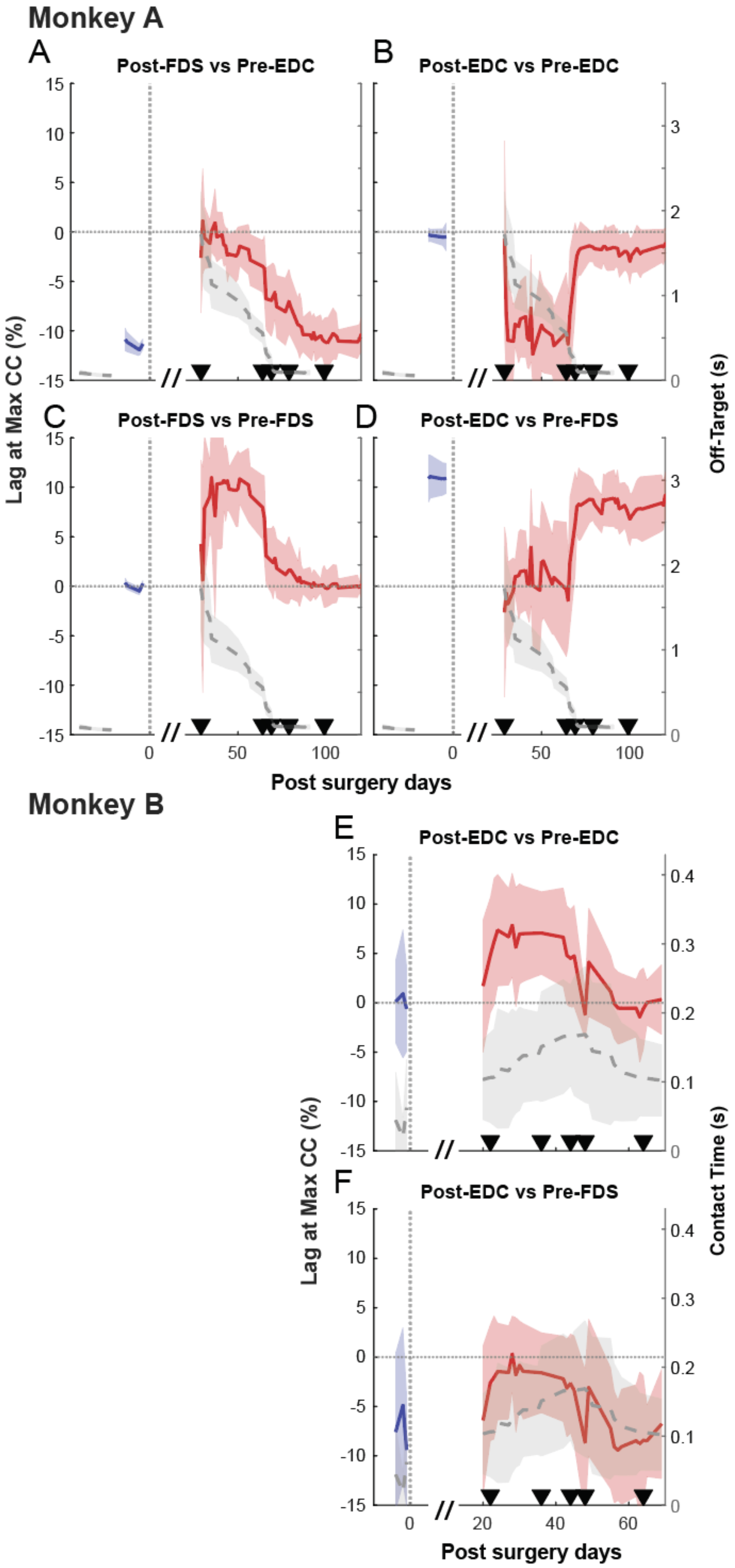
Monkey A
A
Post-FDS vs Pre-EDC
B
Post-EDC vs Pre-EDC
C
Post-FDS vs Pre-FDS
D
Post-EDC vs Pre-FDS
Lag at Max CC (%)
Off-Target (s)
Post surgery days
Monkey B
E
Post-EDC vs Pre-EDC
F
Post-EDC vs Pre-FDS
Lag at Max CC (%)
Contact Time (s)
Post surgery days

91

**Figure S1: Evolution of Time Lag at Peak Cross-Correlation Between Transferred Muscle EMGs.**

Quantification of the optimal time lag (in % task range) yielding the maximum cross-correlation between individual muscle EMG profiles. (A-D) Monkey A; (E-F) Monkey B. Each panel plots the time lag calculated between the recorded EMG profile and the average pre-TT baseline (specific comparisons, e.g., "Post-EDC vs. Pre-EDC," are indicated in titles). Left Y-axis: Time Lag. Blue traces: Pre-surgery data; Red traces: Post-surgery data. Right Y-axis (Gray dashed traces): Behavioral error metrics (Off-target reaching duration for Monkey A; Contact time for Monkey B). A positive lag value indicates that the muscle activity is delayed relative to the pre-surgery baseline; a negative value indicates it is advanced (occurs earlier). The dotted horizontal line indicates zero lag. Shaded envelopes represent standard deviations. Black triangles indicate landmark days.

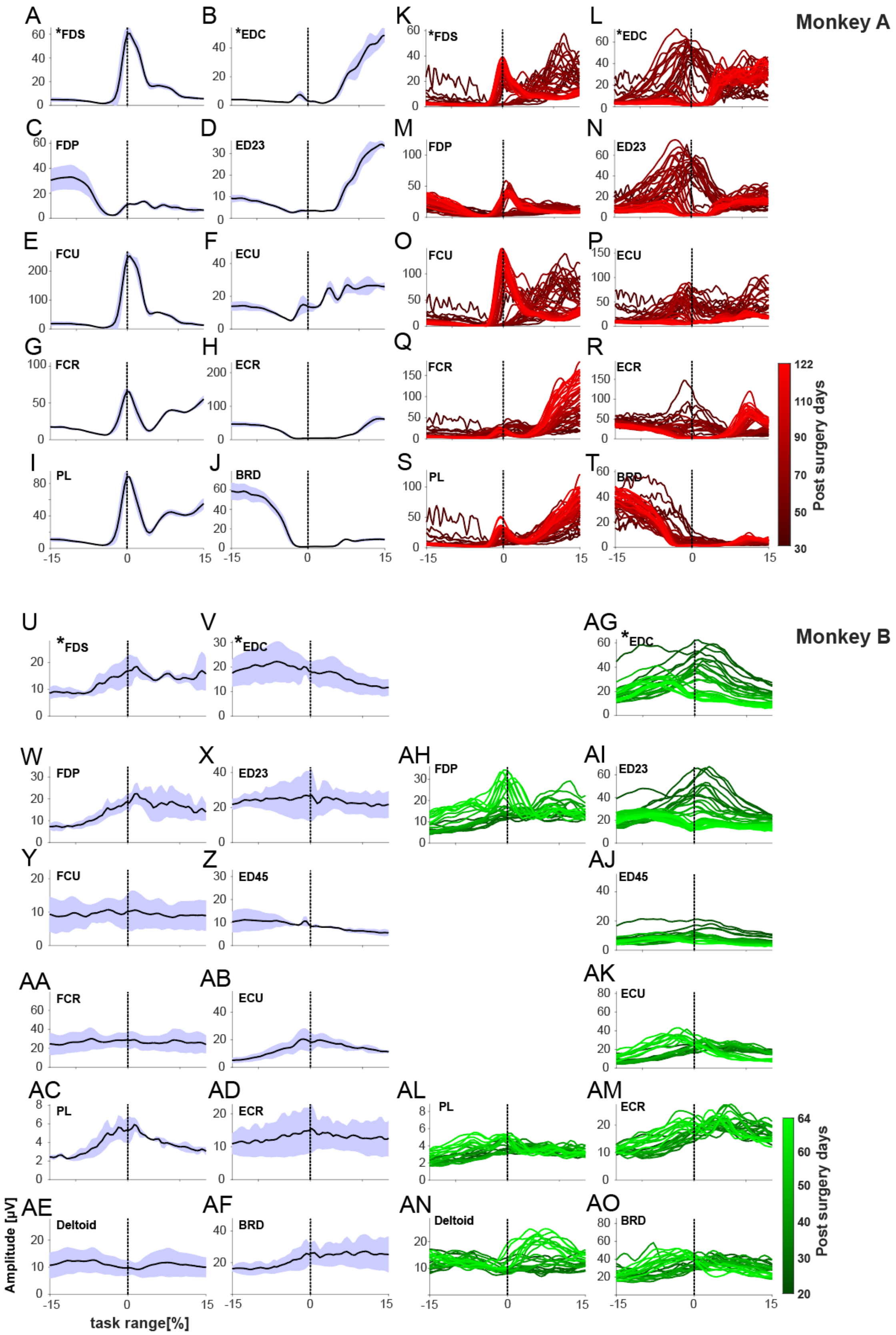
Monkey A
Monkey B
A *FDS
B *EDC
C FDP
D ED23
E FCU
F ECU
G FCR
H ECR
I PL
J BRD
K *FDS
L *EDC
M FDP
N ED23
O FCU
P ECU
Q FCR
R ECR
S PL
T BRD
U *FDS
V *EDC
W FDP
X ED23
Y FCU
Z ED45
AA FCR
AB ECU
AC PL
AD ECR
AE Deltoid
AF BRD
AG *EDC
AH FDP
AI ED23
AJ ED45
AK ECU
AL PL
AM ECR
AN Deltoid
AO BRD
Post surgery days
Amplitude [μV]
task range[%]

**Figure S2: EMG activity profiles for all recorded muscles across all sessions.**

EMG activity profiles [μV] for all recorded muscles in Monkey A (A-T) and Monkey B (U-AO). (A-J, U-AF) Average pre-TT profiles (±SD in blue). (K-T, AG-AO) Post-surgery profiles for each recording day, aligned on object release (Monkey A) or food touch (Monkey B). Lighter colors indicate later recording days (see color bar). Muscles with surgically transferred tendons are marked with an asterisk (*).

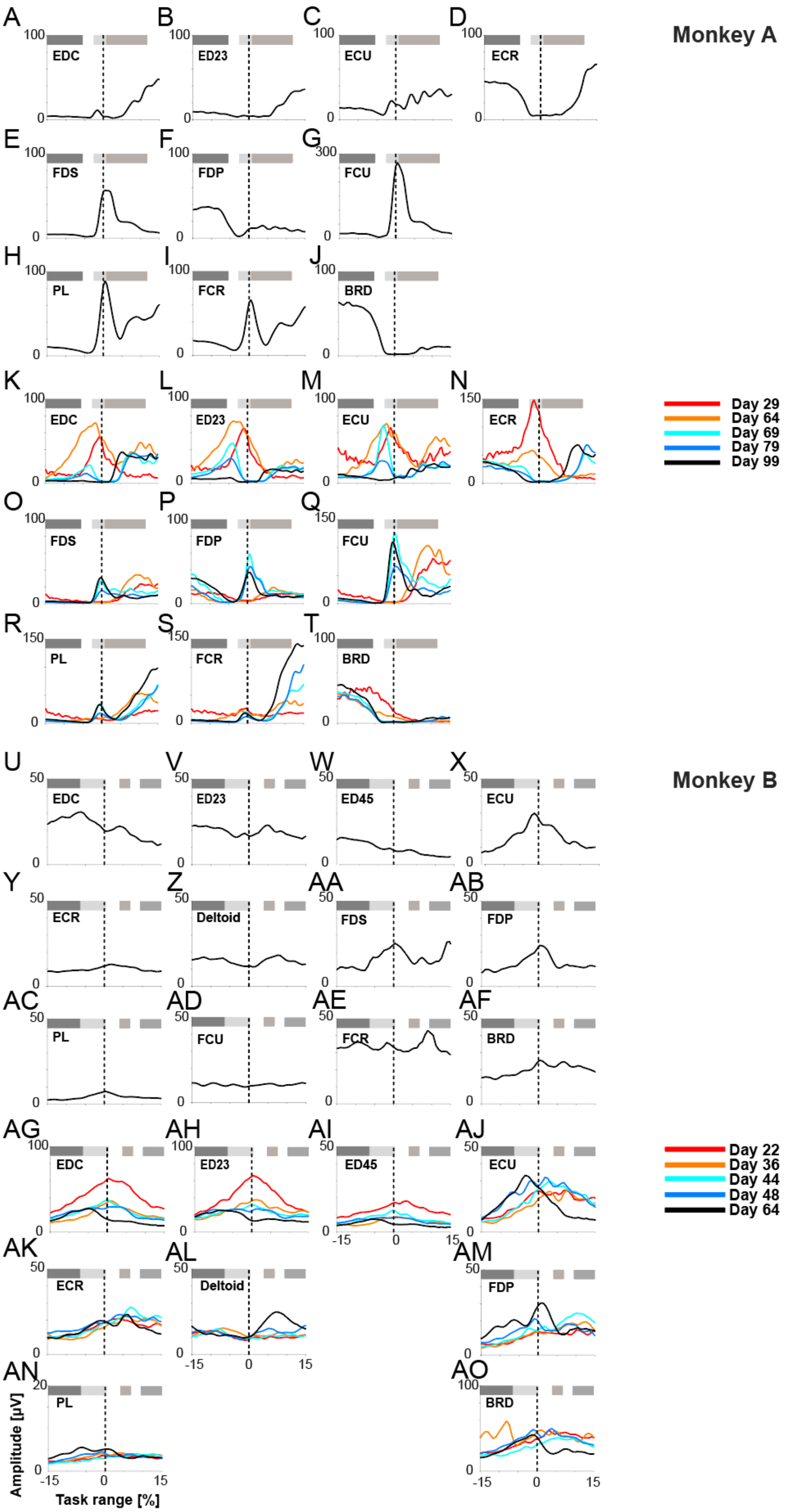
Monkey A
EDC
ED23
ECU
ECR
FDS
FDP
FCU
PL
FCR
BRD
Day 29
Day 64
Day 69
Day 79
Day 99
Monkey B
EDC
ED23
ED45
ECU
ECR
Deltoid
FDS
FDP
PL
FCU
FCR
BRD
Day 22
Day 36
Day 44
Day 48
Day 64
Amplitude [μV]
Task range [%]

**Figure S3: EMG activity profiles of ‘Landmark days’.**

Monkey A: EMG activity profiles [μV] of selected ‘Landmark days’ for all recorded muscles (pre- and post TT, A-J and K-T, respectively). Profiles are aligned on hold off-set (object 1) at zero percent task range [%] (dashed lines). Horizontal gray bars mark the three behavioral periods (hook, grasp and release, respectively). Monkey B: pre- and post TT, U-AF and AG-AO, respectively. Profiles are aligned on LED on-set (food touch) at zero percent task range [%] (dashed lines). Horizontal gray bars mark the four behavioral periods (finger extension, food touch, food pick-up and food transport, respectively).

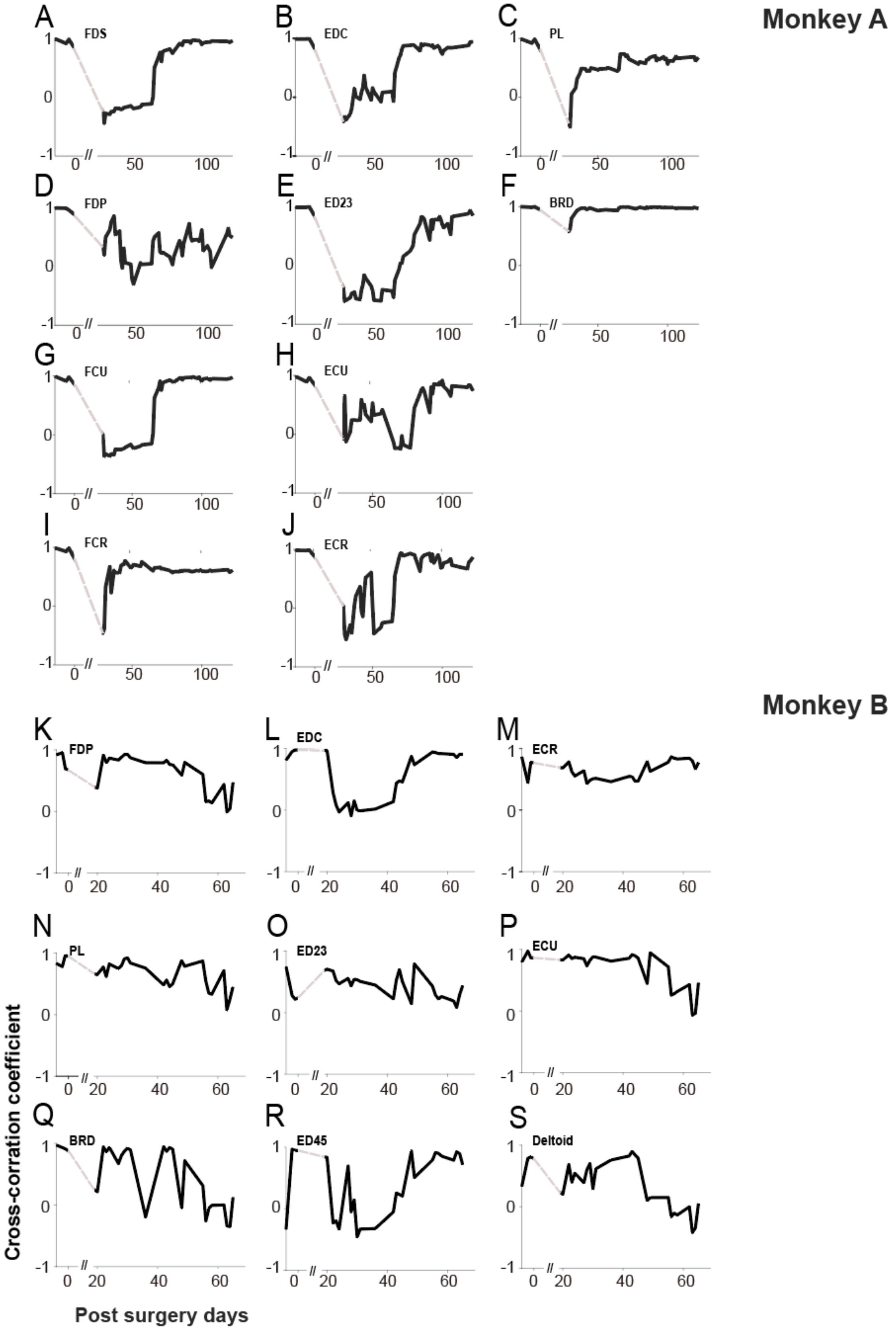
Monkey A
Monkey B
A FDS
B EDC
C PL
D FDP
E ED23
F BRD
G FCU
H ECU
I FCR
J ECR
K FDP
L EDC
M ECR
N PL
O ED23
P ECU
Q BRD
R ED45
S Deltoid
Cross-corration coefficient
Post surgery days

**Figure S4: Cross-correlation analysis for individual muscles.**

Cross-correlation of each muscle's EMG activity profile against its own pre-TT control data. The plots show the evolution of the correlation coefficient over post-surgery days for Monkey A (A-J) and Monkey B (K-S), quantitatively illustrating the time course of adaptation for each muscle.

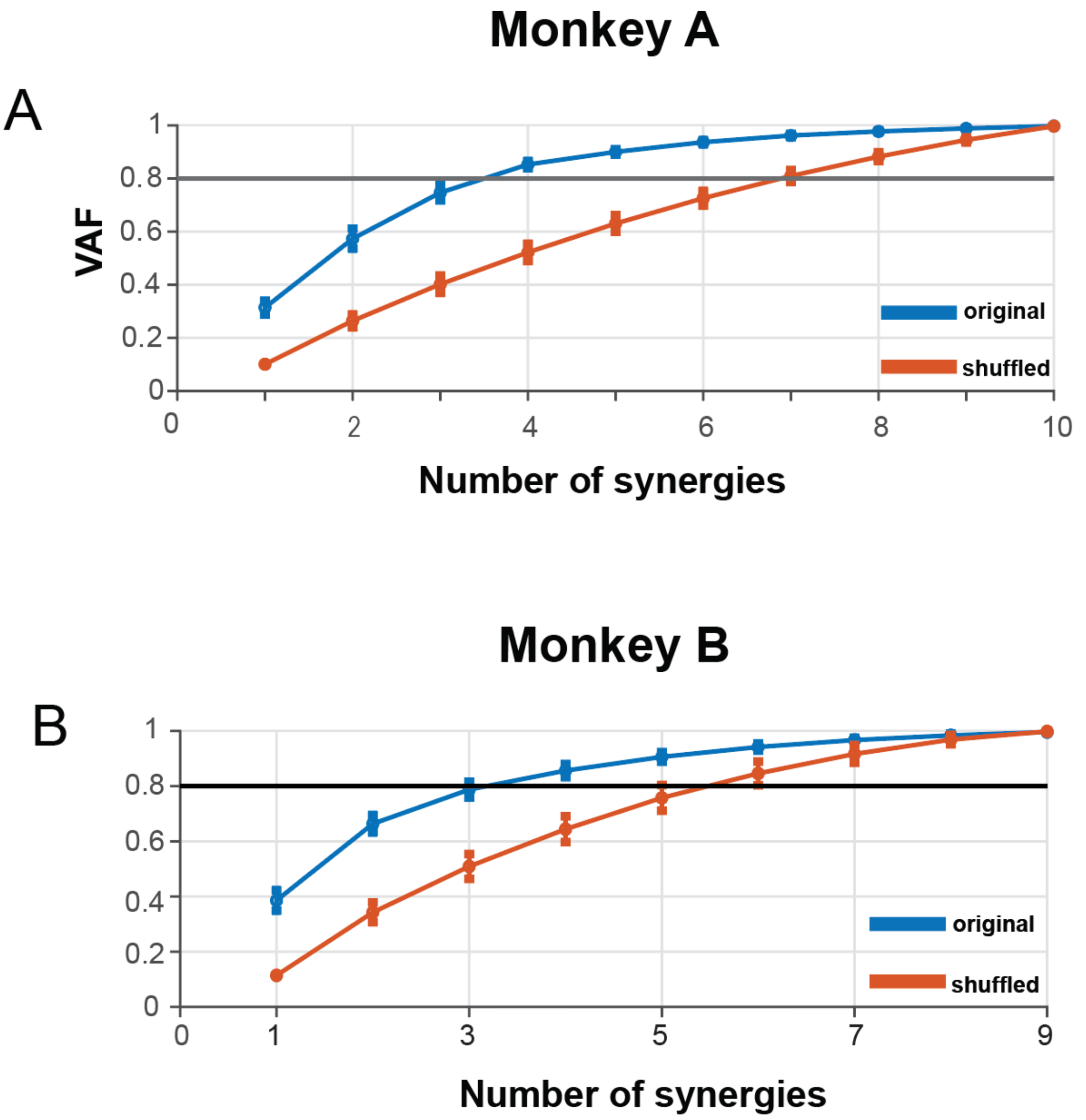


**Figure S5: Variance Accounted For (VAF)**

(A-B) Cumulative Variance Accounted For (VAF) plotted against the number of synergies for Monkey A (bottom) and Monkey B (top). Blue lines: original data; Red lines: shuffled data. The black horizontal line indicates the 80% threshold used to determine the number of synergies.

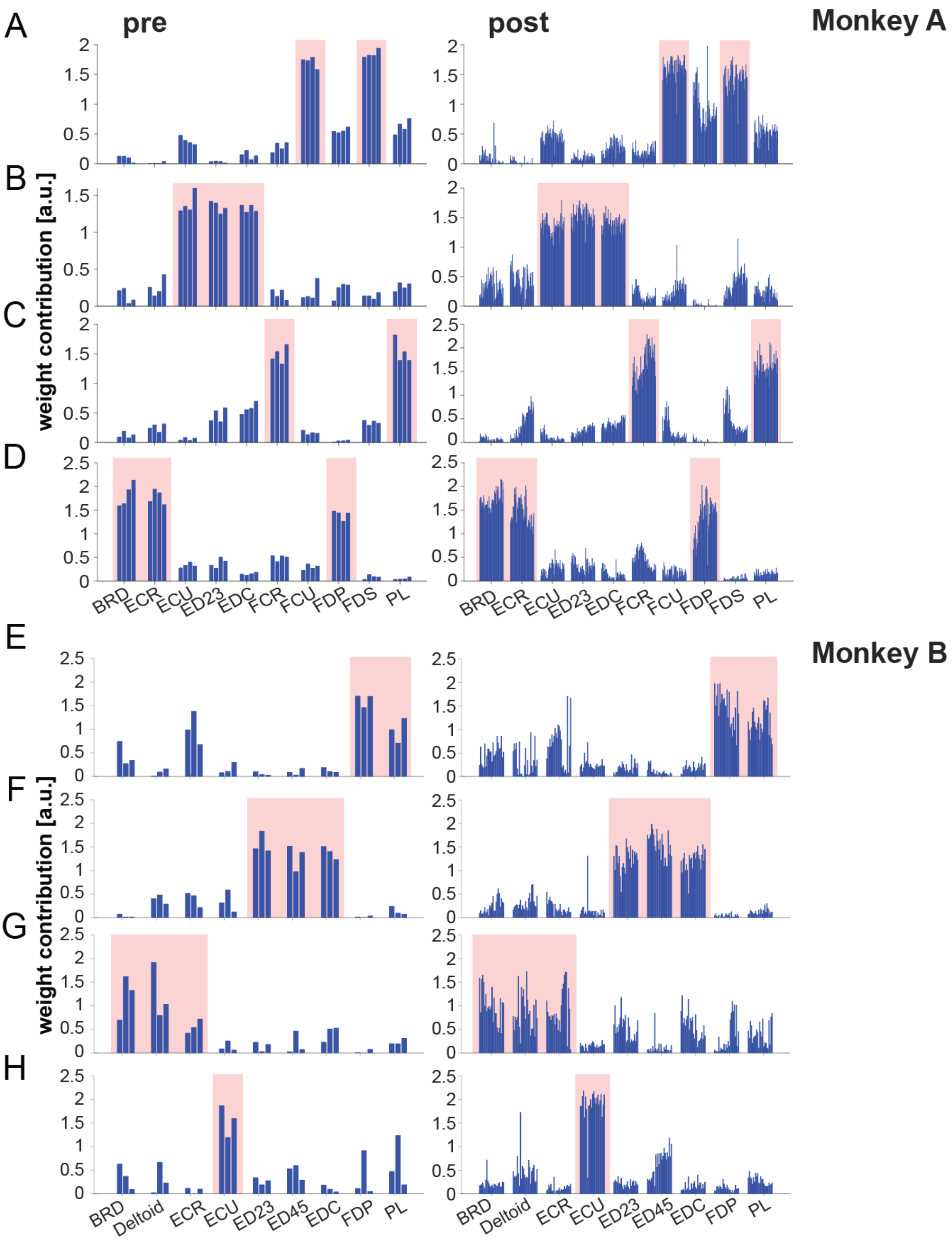


**Figure S6: Synergy weights of all experimental sessions.**

(A-D, monkey A) Synergy weights for each extracted muscle synergy. Each recording day was plotted individually before (left column) and after TT surgery (right column). Red boxes indicate the main contributing muscles to each of the corresponding synergy. (E-F, monkey B). (A, E) Synergy A, (B, F) Synergy B, (C, G) Synergy C and (D, H) Synergy D.

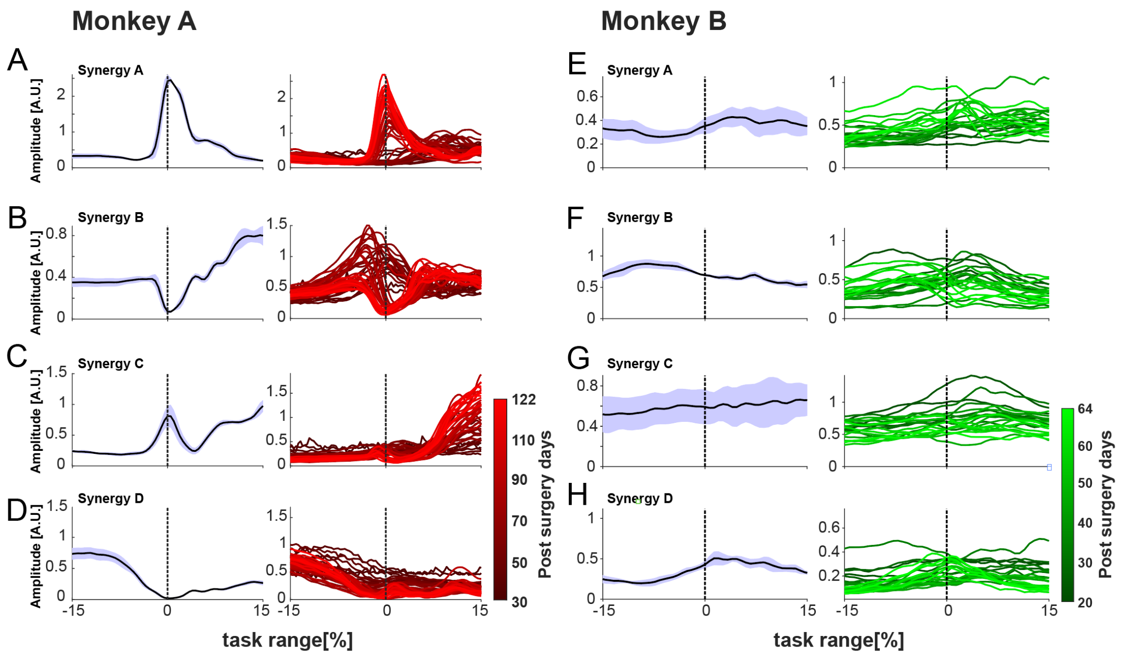


**Figure S7: Time varying activation profiles of muscle synergies.**

(A-D) temporal synergies of monkey A. The left column shows the mean activation profiles of the control data for all four synergies (synergy A-D) with the corresponding standard deviation (blue envelope) aligned on hold off-set (object 1) at zero percent task range [%] (dashed lines). The right column shows the temporal activation profiles for each recording day post-TT surgery (red lines, with darker and lighter colors representing early and late recording days, respectively). (E-H) same as above for monkey B). Profiles are aligned on LED on-set (food touch) at zero percent task range [%].

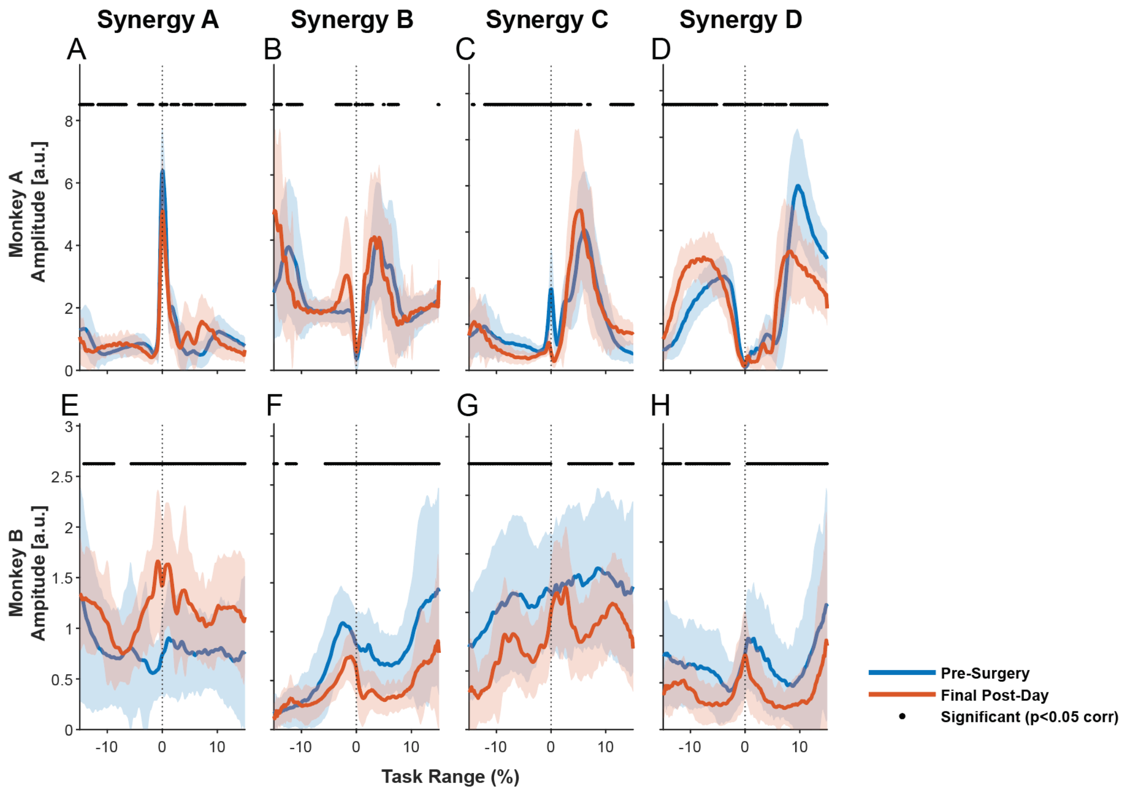


**Figure S8: Quantitative comparison of pre-surgery and final post-surgery synergy activation profiles.**

Comparison of temporal activation profiles (y-axis: Amplitude [a.u.]) for muscle synergies (A–D) between the pooled pre-surgery control period (blue traces; Mean ± SD) and the final post-surgery recording day (red traces; Mean ± SD). (Top Row) Monkey A. (Bottom Row) Monkey B. To identify specific phases where the recovered motor plan differed from baseline, a point-by-point Wilcoxon rank-sum test was performed (Bonferroni-corrected for multiple comparisons). Black markers at the top of each panel indicate time points where the activation amplitude remained statistically significantly different ($p < 0.05$) from the pre-surgery baseline. The widespread presence of these significant differences, even where temporal shapes appear similar, confirms that the final motor state is a functional approximation ('good enough') rather than a perfect restoration of the original control strategy.

207

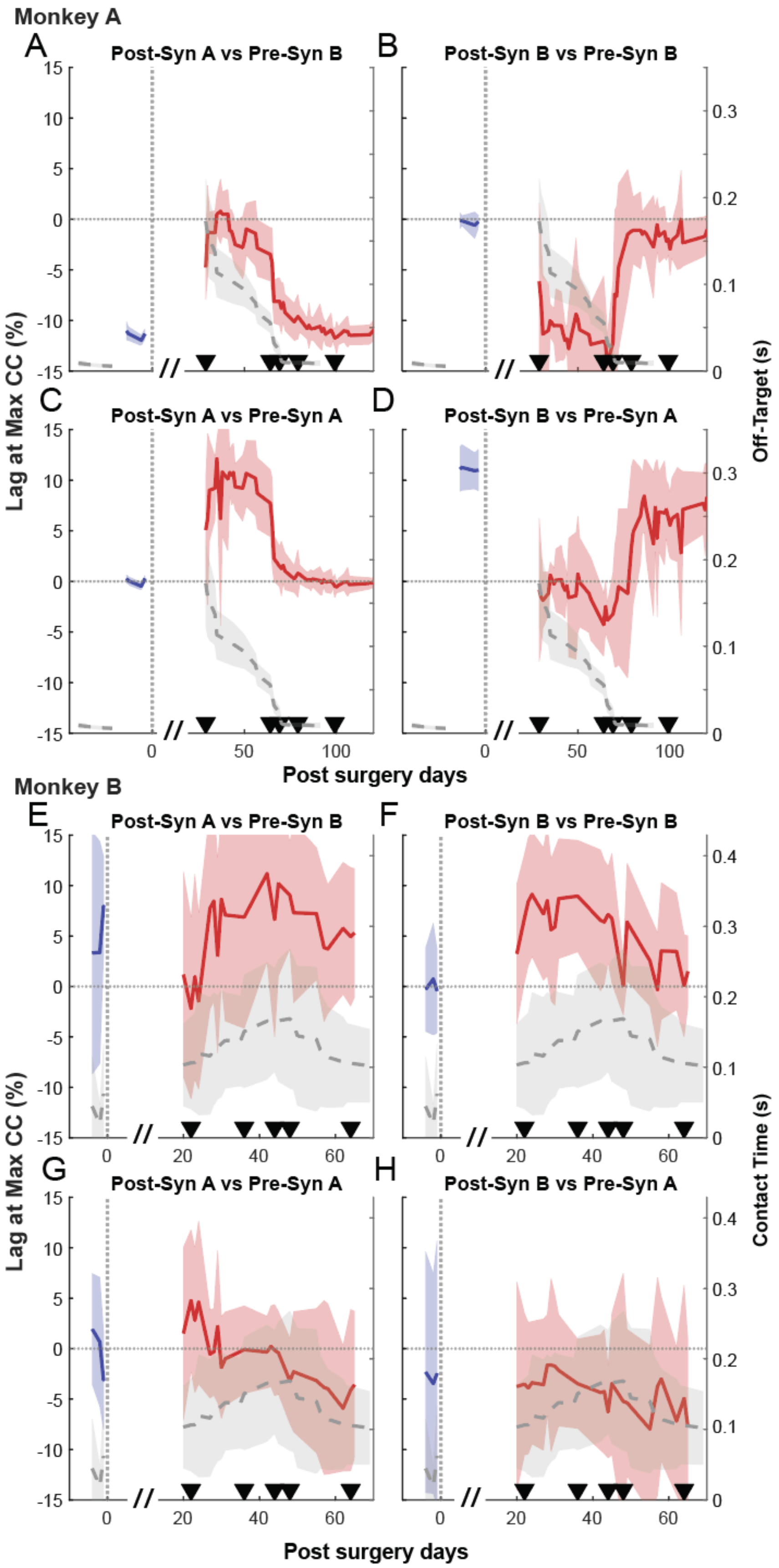
Monkey A
A
Post-Syn A vs Pre-Syn B
B
Post-Syn B vs Pre-Syn B
C
Post-Syn A vs Pre-Syn A
D
Post-Syn B vs Pre-Syn A
Lag at Max CC (%)
Off-Target (s)
Post surgery days
Monkey B
E
Post-Syn A vs Pre-Syn B
F
Post-Syn B vs Pre-Syn B
G
Post-Syn A vs Pre-Syn A
H
Post-Syn B vs Pre-Syn A
Lag at Max CC (%)
Contact Time (s)
Post surgery days

**Figure S9: Evolution of Time Lag at Peak Cross-Correlation Between Primary Synergies.**

Quantification of the optimal time lag yielding the maximum cross-correlation between the Primary Flexor (Synergy A) and Extensor (Synergy B) synergy activation profiles. (A-D) Monkey A; (E-H) Monkey B. Plots show the lag between the recorded synergy activation and the pre-TT baseline (specific comparisons indicated in titles). Left Y-axis: Time Lag. Blue traces: Pre-surgery data; Red traces: Post-surgery data. Right Y-axis (Gray dashed traces): Behavioral error metrics (Off-target reaching duration for Monkey A; Contact time for Monkey B). Positive values indicate the synergy activation is delayed relative to the baseline; negative values indicate it is advanced. Note the temporal correspondence between high behavioral cost (peaks in gray traces) and large fluctuations in synergy timing during the early adaptation phase. Shaded envelopes represent standard deviations.

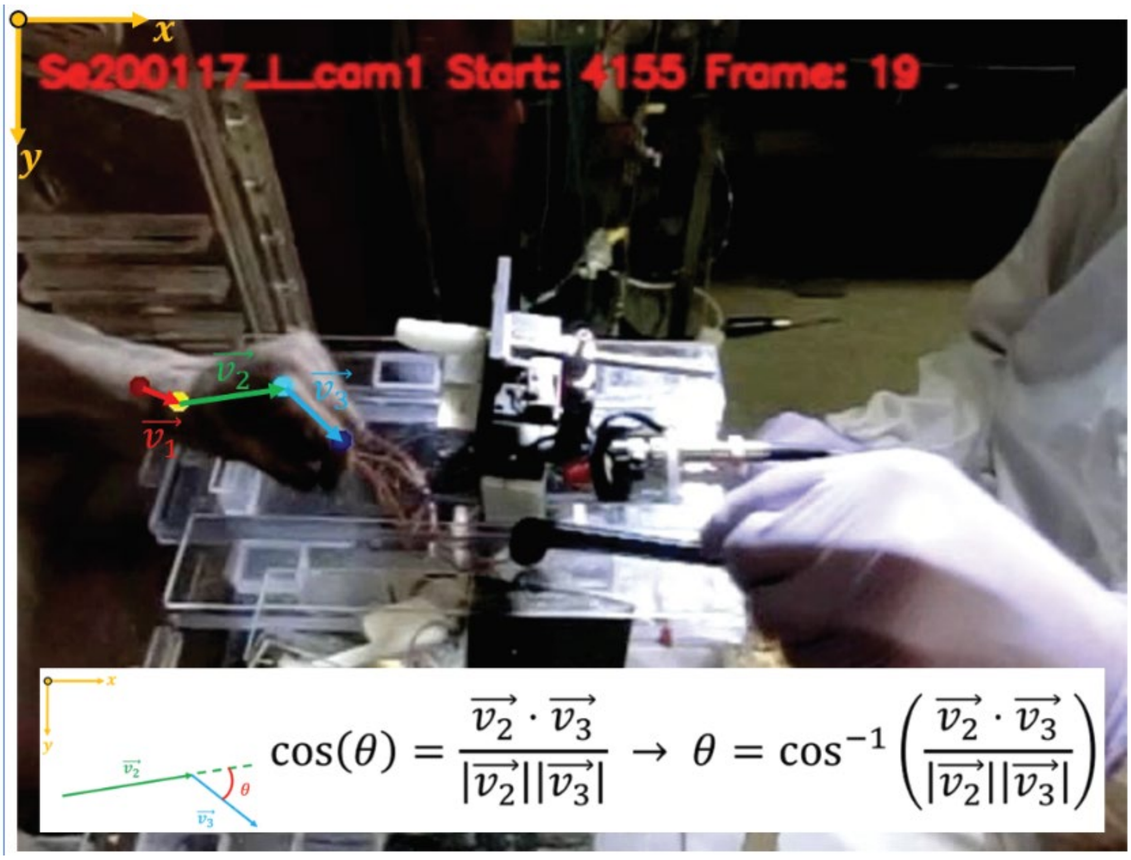


**Figure S10: Joint angle measurement.**

This figure shows an illustration of joint angle measurement using a still picture of the monkey while performing the task. See methods section for further details.

### Supplementary Table 1: Muscle Abbreviations, Functions, and Synergy Groupings

This table provides a reference for all muscles recorded in the study. Columns show the muscle abbreviation, full name, primary functional group, and the main synergy (A, B, C, or D) to which each muscle was assigned for Monkey A and Monkey B.

| Abbreviation | Full Muscle Name | Primary Function Group | Synergy (Monkey A) | Synergy (Monkey B) |
|---|---|---|---|---|
| FCU | Flexor Carpi Ulnaris | Wrist Flexor/Ulnar Deviator | A | N/A[1] |
| **FDS** | **Flexor Digitorum Superficialis** | **Finger Flexor** | **A** | **N/A[1]** |
| ECU | Extensor Carpi Ulnaris | Wrist Extensor/Ulnar Deviator | B | D |
| **EDC** | **Extensor Digitorum Communis** | **Finger Extensor** | **B** | **B** |
| ED2,3 | Extensor Digitorum-2,3 | Finger Extensor | B | B |
| FCR | Flexor Carpi Radialis | Wrist Flexor/Radial Deviator | C | N/A[1] |
| PL | Palmaris Longus | Wrist Flexor | C | A |
| BRD | Brachioradialis | Elbow Flexor/Wrist Radial Deviator | D | C |
| ECR | Extensor Carpi Radialis | Wrist Extensor/Radial Deviator | D | C |
| FDP | Flexor Digitorum Profundus | Finger Flexor | D | A |
| DEL | Deltoid | Shoulder Abductor/Flexor | N/A | C |
| ED4,5 | Extensor Digitorum-4,5 | Finger Extensor | N/A | B |

***Bolded muscles (EDC and FDS) were the targets of the tendon transfer surgery.***

***([1])****: EMG signal for this muscle was lost post-surgery in Monkey B and was excluded from the synergy analysis.* ***(N/A)****: Muscle not recorded or included in the analysis for that monkey.*

### Supplementary Table 2:

Statistical comparison of Pre-Surgery vs. Final Post-Surgery Synergy Profiles

| Monkey | Synergy | Cosine Similarity (Shape) | Wilcoxon P-value (Global Amplitude) | Permutation Test P-value (Trajectory) |
|---|---|---|---|---|
| **Monkey A** | **A** | 0.9243 | 0.3451 (n.s.) | < 0.0001* |
| | **B** | 0.9483 | **0.0019*** | < 0.0001* |
| | C | 0.9163 | 0.3808 (n.s.) | < 0.0001* |
| | D | 0.9433 | 0.3541 (n.s.) | < 0.0001* |
| **Monkey B** | **A** | 0.9043 | **< 0.0001*** | < 0.0001* |
| | **B** | 0.9726 | **< 0.0001*** | < 0.0001* |
| | C | 0.915 | **< 0.0001*** | < 0.0001* |
| | D | 0.9017 | **< 0.0001*** | < 0.0001* |

*(Note: "**Global Amplitude**" refers to a comparison of the distribution of mean activation values across the entire task cycle. "**n.s.**" = not significant).*
